# From expansion to consolidation: two decades of Gene Ontology evolution

**DOI:** 10.64898/2026.03.04.709507

**Authors:** Borja Pitarch, Florencio Pazos, Monica Chagoyen

## Abstract

The Gene Ontology (GO) is a long-standing, community-maintained knowledge resource that underpins the functional annotation of gene products across numerous biological databases. Released regularly, GO and its associated annotations form a large, continuously evolving dataset whose temporal dynamics have direct consequences for data reuse, versioning, and reproducibility. Because analytical results derived from GO are inherently tied to specific ontology and annotation releases, a systematic understanding of how GO changes over time is essential for transparent interpretation and long-term reuse of GO-based analyses. Here, we present a comprehensive temporal characterization of the Gene Ontology and its annotations spanning 21 years of publicly available releases. Treating successive ontology and annotation versions as longitudinal research data, we quantify changes in ontology structure, term composition, relationships, and annotation content across time and across three representative annotation resources. Our analysis reveals sustained growth of GO over its lifetime, accompanied by marked structural reorganization, particularly affecting high-level, general ontology terms. Notably, across multiple structural and annotation metrics, we identify a transition toward increased stability beginning around 2017, consistent with a maturation phase of the resource. This work provides a reference framework for researchers who rely on GO releases for data integration, benchmarking, and reproducible functional analysis.

## Introduction

The Gene Ontology (GO) is the most widely used ontology in molecular biology and a core community standard for the functional annotation of gene products across a broad range of organisms (Gene Ontology Consortium 2026). Distributed through regular public releases, GO provides a structured and computable vocabulary that underpins the annotation, integration and reuse of genome-scale and other high-throughput omics datasets. As such, GO and its associated annotations constitute a foundational research data resource whose properties directly affect the interpretation, comparability, and reproducibility of a wide range of downstream analyses.

From its inception, GO was conceived as a dynamic and continuously evolving resource, designed to reflect the current state of biological knowledge (Ashburner *et al*. 2000). Since the establishment of the GO Consortium in 1998 and the first formal description of the ontology in 2000, ontology development and annotation have been carried out through a distributed, community-driven curation process. Ontology updates are released regularly and may involve minor, localized changes to individual terms or more extensive revisions affecting entire ontology branches. GO is organized into three subontologies—Biological Process (BP), Cellular Component (CC), and Molecular Function (MF)—each represented as a directed acyclic graph (DAG) linking the vocabulary terms in a partially hierarchical structure. This structure supports annotation at varying levels of specificity, from general terms at higher levels in the ontology down to highly specific leaf terms, enabling flexible reuse of annotations across diverse data types and analytical contexts.

As biological knowledge expands through new experimental data, publications, and methodological advances, gene annotations must be continuously revised to incorporate newly characterized functions and supporting evidence. At the same time, the underlying ontology must evolve in a controlled and transparent manner, reflecting community consensus, conceptual refinement, and structural consolidation. This coordinated evolution is critical for reproducibility, as gene annotations are released against specific versions of the Gene Ontology (GO), and their interpretation is therefore inseparable from the exact ontology state in which they were generated. Explicit versioning of both the ontology and its annotations ensures traceability over time and enables consistent reanalysis of functional data as GO evolves.

Previous studies have examined aspects of GO evolution over limited time spans or from specific analytical perspectives. Early temporal analyses reported branch-specific patterns of change, with differences in growth and refinement observed across BP, CC, and MF (Dameron, Bettembourg, and Le Meur 2013). Subsequent work demonstrated that structural and annotation changes across successive GO releases can alter the outcomes of commonly used computational analyses, including functional enrichment and similarity measures (Tomczak *et al*. 2018, Paul, Anand, and Pyne 2019, Chen *et al*. 2024). More recent analyses have shown that GO evolution is not purely incremental but can involve punctuated, curator-driven reorganisations, such as the large-scale restructuring of the CC branch in 2019, which substantially modified upper-level ontology organisation while retaining most terms (Valverde *et al*. 2025).

Together, these studies highlight that GO is a long-lived, evolving data resource whose temporal dynamics reflect not only the ongoing accumulation of experimental data but also the expert-driven process of curating and organising a shared vocabulary to represent biological knowledge. For researchers who rely on GO releases for data annotation, integration, benchmarking, or longitudinal analysis, understanding how the ontology and its annotations change over time is essential for transparent reuse and reproducibility. However, a comprehensive, long-term characterisation of GO evolution that treats successive releases as a unified longitudinal dataset, including both ontology and annotations, remains limited.

In this work, we present a large-scale temporal analysis of the Gene Ontology and its annotations spanning 21 years of public releases (2004–2024). By systematically characterising changes in ontology structure and annotation content across time, we provide a data-driven description of GO evolution that supports informed reuse of this widely adopted resource and contributes to broader discussions on managing evolving ontologies as shared research data.

## Results

The three subontologies of the Gene Ontology (GO) —Biological Process (BP), Molecular Function (MF), and Cellular Component (CC)— represent different aspects of gene function, so we analyzed them separately. Treating each subontology independently allows us to characterise structural and annotation changes specific to the type of knowledge captured, as well as to highlight differences in how these data are maintained, updated, and reused. Such separation is particularly relevant for understanding the temporal properties of GO releases, since the subontologies follow different patterns of term growth, relationship refinement, and annotation density, which can influence downstream computational analyses and reproducibility of results.

Our analysis is structured in two parts. In the first, we focus on the ontology itself, examining the number of terms, relationships, and overall DAG structure across successive releases. In the second, we examine a representative set of annotation datasets associated with GO, evaluating their growth, coverage, and temporal dynamics as longitudinal data resources. This approach provides a comprehensive view of GO as an evolving dataset, encompassing both its conceptual framework and the annotations that operationalise it for data integration and reuse.

### Evolution of the ontology

Each GO release contains two types of terms: active terms, which are currently used for annotation, and obsolete terms, which have been deprecated in previous releases. Obsolete terms are retained to preserve historical context and ensure traceability, but they should not be assigned to gene products in new annotations.

Although the sizes of the subontologies differ substantially (26,467 terms in GO:BP, 10,146 in GO:MF, and 4,022 in GO:CC, according to the latest release analyzed), all show a similar temporal trend. In each subontology, the total number of active GO terms increased steadily until approximately 2017, after which it plateaued (Figure 1a). A slight decrease is observed for BP and CC over the subsequent 5–6 years, whereas MF shows a more pronounced decrease in 2024. Notably, the rate of increase is similar across the three ontological aspects during the early years (up to 2011), with modest divergence thereafter, particularly between BP and MF in the following period, until growth slowed and eventually reversed around 2017.

**Figure 1:**
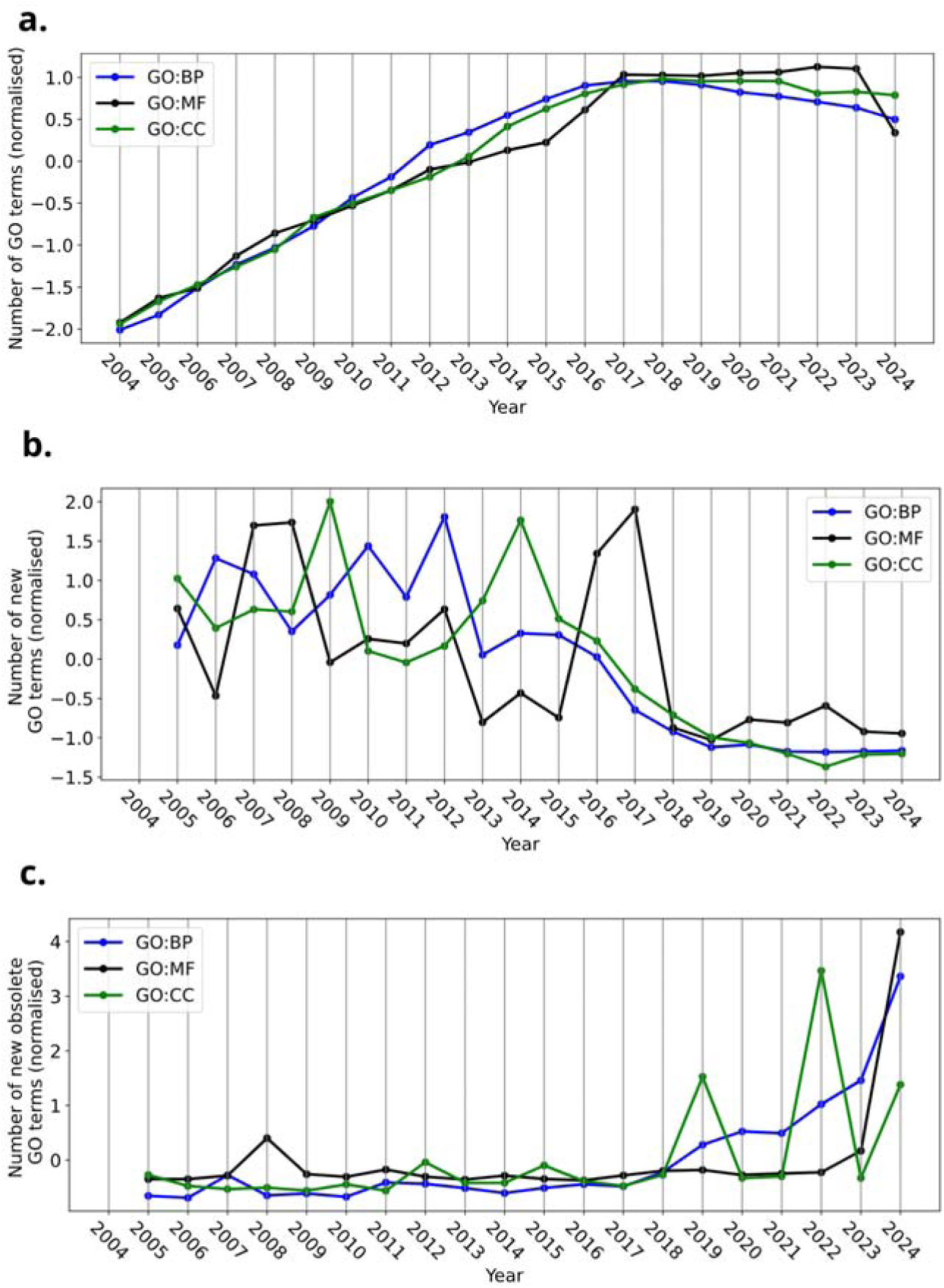
Evolution of GO terms by subontology. a) Number of active GO terms by year. b) Evolution of the number of new GO terms added each year. c) Number of terms obsoleted terms each year.

The net change of term counts reflects a balance between two processes: the addition of new terms and the obsoletion of existing ones. We therefore analyzed these factors separately. The rate of new term additions declined in all three subontologies beginning around 2017 (Figure 1b). In parallel, the number of newly obsoleted terms began to increase for BP and CC, with greater variability in CC, whereas for GO:MF this increase occurred more recently, starting in 2023 (Figure 1c).

Most terms exhibit lifespans of 12 years or longer, and a substantial fraction remain unchanged throughout the entire 21-year study period (Figure S1). Among the three subontologies, GO:MF contains the highest proportion of long-lived, stable terms.

To highlight the biological topics emphasized by GO at different time points, we performed a word enrichment analysis on newly introduced terms for each year (see Methods). Figure 2 illustrates examples from three representative years: 2009 shows enrichment for gland-related terms, 2010 for kidney-related terms, and 2016 for highly significant nervous system–related additions. Results for all other years are provided in Figures S2 to S19.

**Figure 2:**
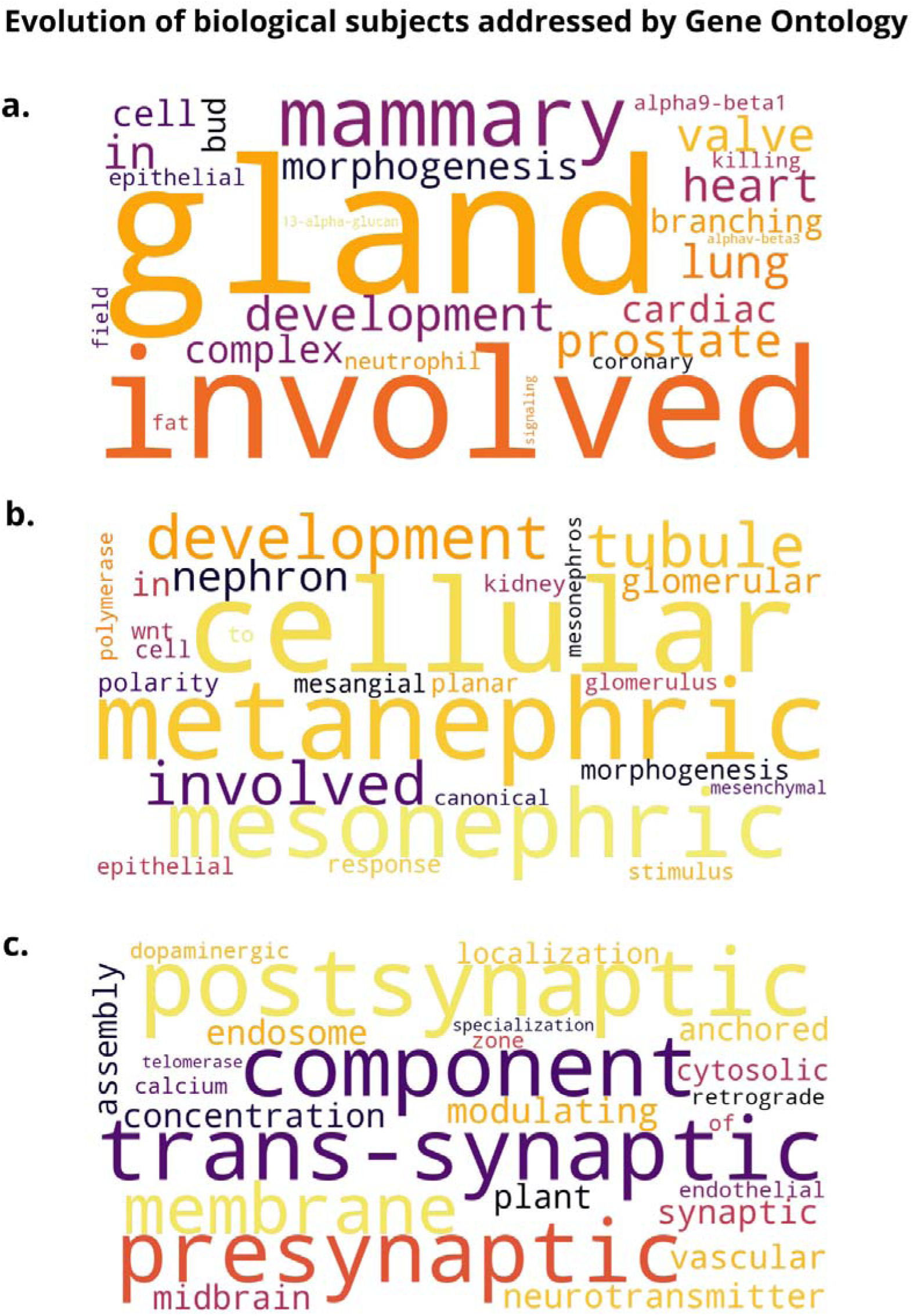
Words enriched in names and synonyms of GO terms added in a) 2009, b) 2010 and c) 2016.

Because GO represents biological knowledge at multiple levels of specificity, temporal changes are expected to affect its structure. Since gene annotations are generally made at the most specific level possible, we classified GO terms by their position in the DAG into two categories: *leaf terms*, representing the most specific concepts, and *non-leaf terms*, encompassing more general terms. To characterise structural changes over time, we examined the ratio of non-leaf to leaf terms for each subontology (Figure 3a). GO:MF consistently exhibits the highest proportion of leaf terms, followed by GO:CC. In contrast, GO:BP is the only subontology in which non-leaf terms consistently outnumber leaf terms (ratio > 1.0), reflecting a more complex hierarchical organisation. This complexity increases until approximately 2018, after which it declines and stabilizes. In light of the overall term growth trends (Figure 1), these results suggest that the GO:BP expansion prior to 2017 was driven primarily by the addition of internal (non-leaf) terms, whereas GO:CC and GO:MF remained comparatively stable in this respect.

**Figure 3:**
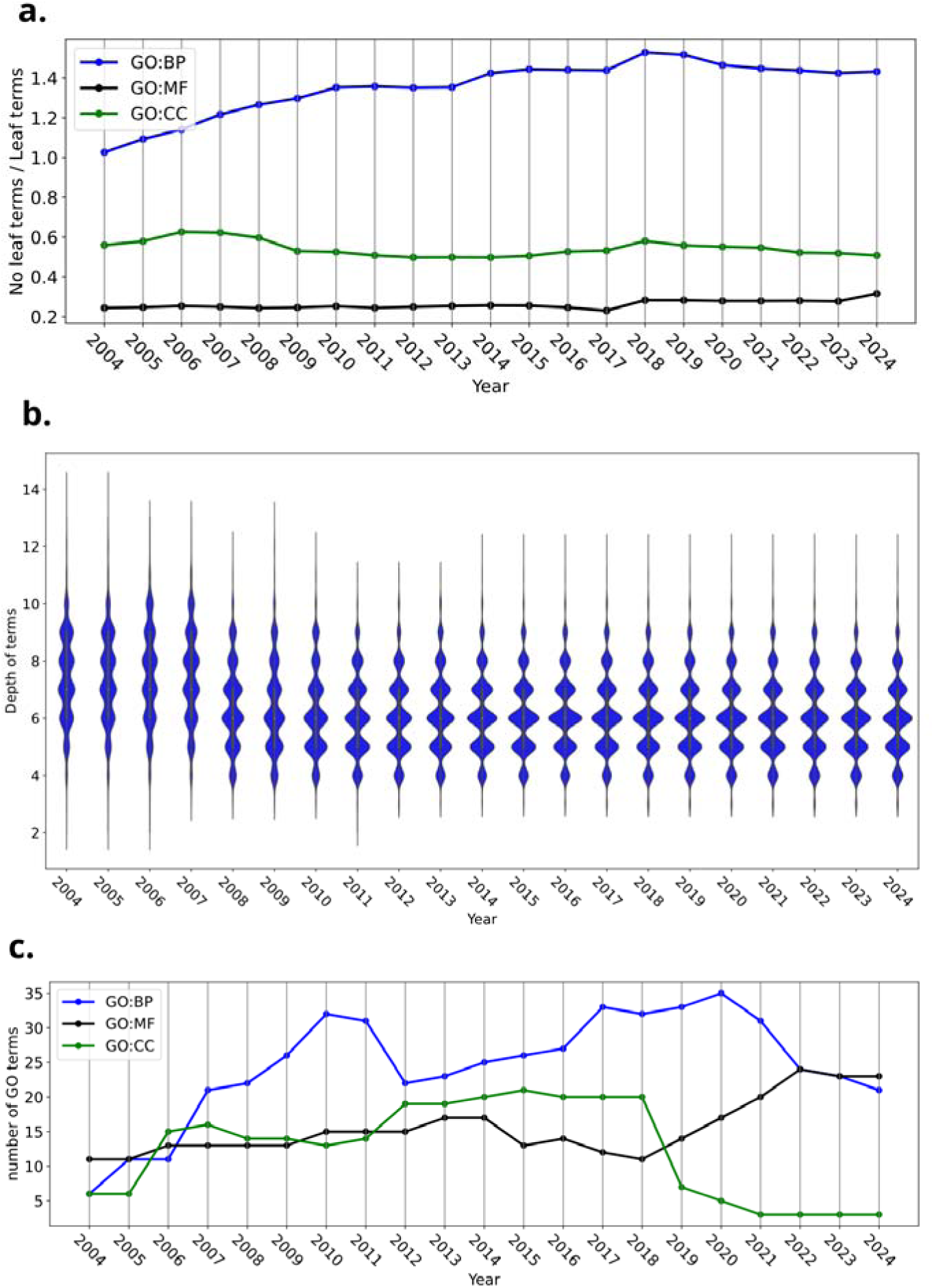
Evolution of GO structure by subontology. a) Ratio of non-leaf terms per leaf term in GO per year. b) Distribution of GO:BP term depth per year. c) Number of first layer terms per year

The addition of internal nodes is expected to influence term depth, defined as the distance from the root of the ontology. Figure 3b shows the distribution of term depths for GO:BP over time. Surprisingly, median depths decreased until around 2011, indicating that new internal terms tend to expand existing branches laterally (broadening the ontology) rather than extending paths linearly away from the root.

The first layer of the ontology, comprising the broadest and least specific terms, provides the foundational classification within each subontology. Figure 3c shows the size of this layer over time for all three subontologies. Large fluctuations in the first layer reflect major structural reorganisations affecting the ontology’s overall framework. Since 2017, the size of the first layer has decreased for GO:BP and GO:CC, increased slightly for GO:MF, and remained relatively stable from 2022 onward.

GO structure is defined by typed parent-child relationships among terms. To analyze their temporal evolution, we focused on the most prevalent relationship types (*is_a* and *part_of*) and grouped all remaining types together. Other relationship types were introduced in 2008 for GO:BP and later, in 2018, for GO:MF and GO:CC. Overall, the total number of relationships increased uniformly until 2016 (Figure S20), after which GO:CC exhibited slightly divergent trends, consistent with differences in term dynamics described above. To disentangle these effects, we examined the ratio of relations per terms (Figure 4).

**Figure 4.**
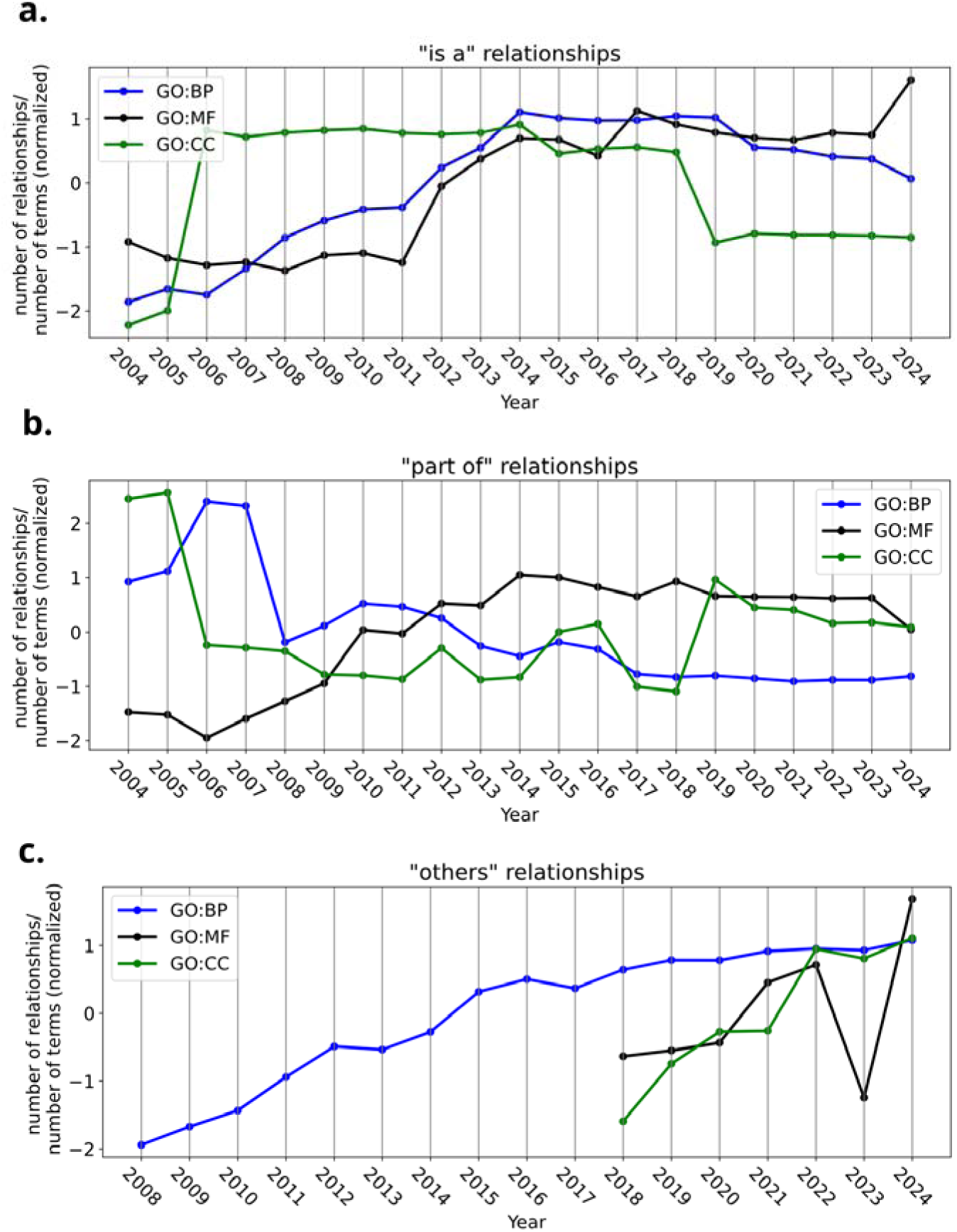
Evolution of GO relations per term by subontology. a) Number of *is_a* relations per term. b) Number of *part_of* relations per term. c) Number of other types of relations.

**Figure 5:**
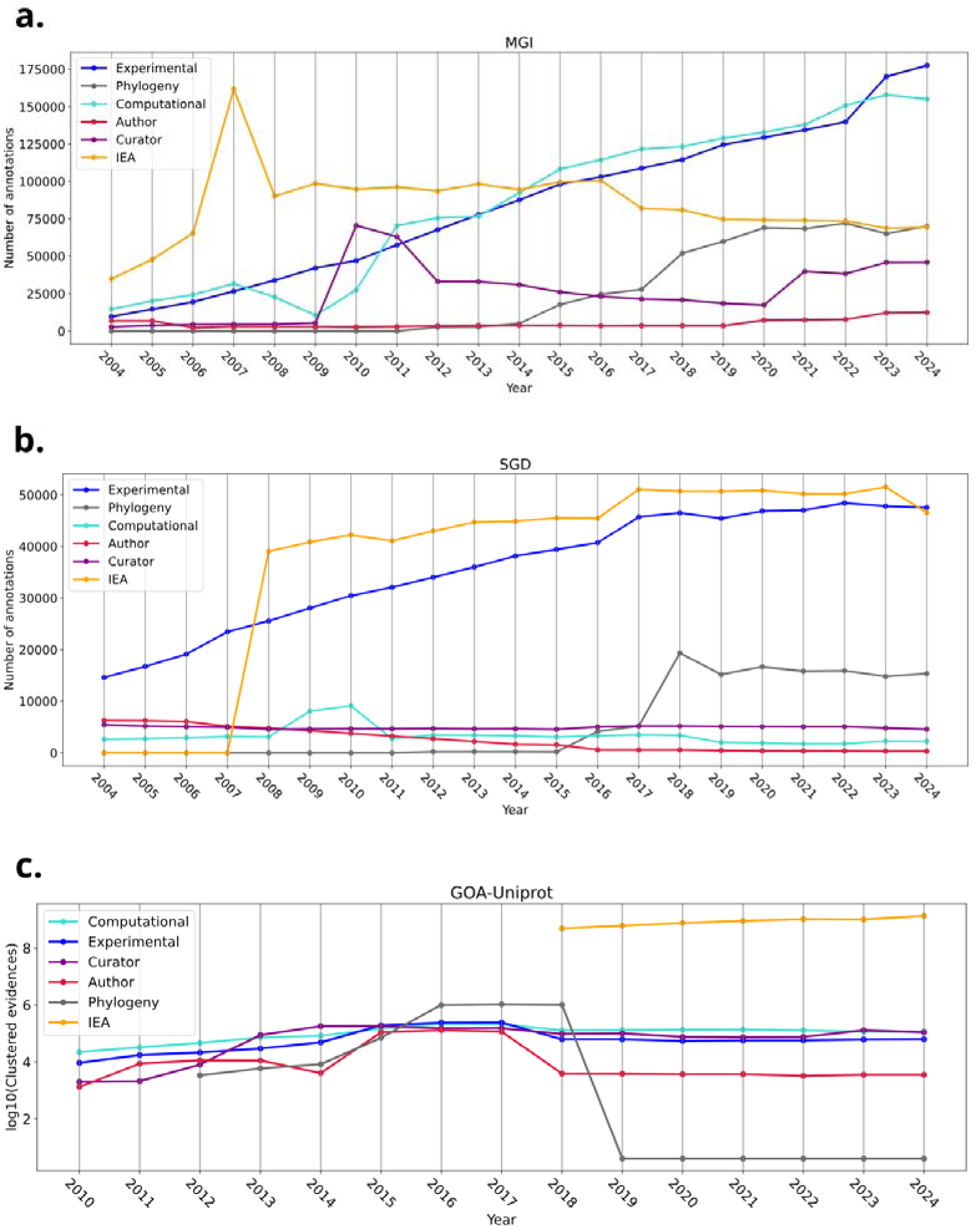
Evolution of the number of annotations per evidence class in a) MGI (*M. musculus*), b) SGD (*S. cerevisiae*) and c) GOA-Uniprot (multi-species).

The ratio of *is_a* relationships per term increased during the first decade analyzed for GO:BP and GO:MF before stabilizing, whereas GO:CC shows a sharp increase in 2006 followed by a decline beginning in 2019. In contrast, the ratio of *part_of* relationships decreased during the early years (with a sharp decrease in 2006 in GO:CC and a relevant increase in the same year, with a pronounced drop in 2006 for GO:CC and a transient increase in GO:BP in the same year, followed by an abrupt decrease in 2008. in GO:BP but followed by an abrupt decrease in 2008). Other relationship types have steadily increased in GO:BP since their introduction in 2008, with similar trends observed more recently in GO:MF and GO:CC. The opposing and abrupt changes observed in *is_a* and *part_of* ratios for GO:CC likely reflect major structural reorganizations of this subontology.

### Evolution of annotations

GO annotations are contributed by multiple member databases, each operating distinct curation and annotation pipelines tailored to their organisms and data sources. Although all contributors adhere to GO Consortium annotation standards and quality-control procedures, differences in curation scope, release schedules, and automated workflows result in heterogeneous annotation dynamics across databases. This variability might be particularly evident for electronically inferred annotations (IEAs), which are produced by database-specific pipelines that combine shared resources—such as InterPro-GO mappings and orthology-based projections—with locally implemented filtering, update, and quality-control strategies. As a result, temporal fluctuations in IEA counts may reflect changes in automated pipelines, curation policies, or upstream mappings, rather than shifts in underlying knowledge alone.

To capture complementary perspectives on annotation dynamics across the GO annotation ecosystem, we performed a temporal analysis of the *Saccharomyces cerevisiae* annotations from SGD (Engel *et al*. 2025), mouse (*Mus musculus*) annotations from MGI (Baldarelli *et al*. 2024), *C. Elegans* annotations from Wormbase (Sternberg *et al*. 2024) and the GOA_Uniprot resource (Huntley *et al*. 2015). *S. cerevisiae* represents a deeply studied model organism with long-standing, intensive manual curation and a high proportion of experimentally supported annotations, providing insight into annotation behaviour in a mature and well-characterised system. Mouse annotations reflect a complex multicellular model with broad biomedical relevance and heterogeneous evidence profiles, illustrating how large-scale experimental programs and integrative curation practices shape annotation evolution over time. GOA, maintained by UniProt, offers a contrasting perspective by aggregating annotations across a wide taxonomic range and combining extensive automated pipelines with targeted manual curation, making it representative of high-throughput, cross-species annotation strategies. Together, these resources enable comparative analysis of annotation growth, evidence composition, and temporal stability across different curation models and research intensities.

The temporal evolution of annotation counts differs across these resources, although all three exhibit an overall increasing trend (Figure S21). In MGI, annotations increase steadily across all GO aspects, with the exception of a local peak in 2007, and with a steeper growth rate observed for GO:BP. In SGD, annotation growth occurs at varying rates: a moderate increase between 2004 and 2007, a pronounced expansion in 2008, and a more gradual increase from 2008 to 2019, followed by stabilization (with a slight decrease in GO:BP during the most recent two years). GOA displays two main phases: a steady increase between 2010 and 2017, a sharp rise in 2018, and a more moderate growth rate from 2019 to 2024.

Given the heterogeneous nature of annotations, we next examined their associated evidence codes, grouped into six broad classes: Experimental, Phylogeny, Computational, Author, Curator and “Inferred from electronic annotation” (IEA). This analysis reveals marked differences between source databases. Both MGI and SGD show a sustained increase in experimentally supported annotations over time, with evidence of stabilization in recent years for yeast, whereas the number of experimental annotations in GOA remains comparatively constant throughout the analyzed period.

Distinct temporal patterns are also observed for IEAs. In MGI, IEA counts peak in 2007, followed by a sharp decline and subsequent stabilization, with a slight decrease after 2016. In SGD, IEAs increase dramatically in 2008, continue to rise modestly in subsequent years, and then remain largely stable. Historical GOA data contain no IEAs prior to 2018; from that point onward, IEA counts remain relatively stable. Finally, we note a pronounced decline in phylogeny-based annotations in GOA beginning in 2019.

## Discussion

The Gene Ontology (GO) has become the de facto standard for the functional annotation of gene products and underpins a wide range of analytical approaches in modern biology. As such, GO plays a central role in many data analysis workflows and is therefore subject to scrutiny from both its developers and the broader scientific community (Pitarch *et al*. 2025). Among the most widely used GO-based methods is gene set enrichment analysis (Reimand *et al*. 2019), which is routinely applied to interpret large gene or protein sets derived from omics experiments. Importantly, the outcomes of enrichment analyses depend directly on the structure and content of GO. Several studies have shown that enrichment results obtained using different temporal versions of GO can differ substantially, in some cases leading to markedly different biological interpretations (Groß *et al*. 2012, Clarke *et al*. 2013, Tomczak *et al*. 2018). Consequently, GO-based results are intrinsically time-dependent, as they are tied to the specific ontology and annotation versions used. This has clear implications for reproducibility and motivates the need to systematically quantify how GO evolves over time and to identify prevailing trends in its development.

GO is an actively maintained resource that undergoes continuous updates, including changes to terms, relationships, and annotations. Our analysis confirms an overall long-term growth trend, with increases in the number of terms and relationships over time. However, between approximately 2017 and 2019, we observe a marked stagnation in the number of terms and certain relationship types, accompanied by an increase in term obsoletion and a reduced rate of new term addition in subsequent years. Most GO terms exhibit considerable stability, with relatively long lifespans, indicating that changes are generally incremental rather than disruptive.

In terms of biological content, the thematic focus of newly added GO terms varies over time, reflecting shifting research priorities and targeted ontology expansions. Several of the enriched topics identified in our analysis correspond to documented GO development efforts, such as the 2016 expansion addressing Parkinson’s disease (Foulger *et al*. 2016), which resulted in the addition of numerous nervous system–related terms. These observations highlight the responsiveness of GO to emerging biomedical research areas and community-driven needs.

At the level of GO’s hierarchical organization, we find that leaf terms—representing the most specific functional concepts—are generally stable over time, with the notable exception of GO:BP, where their number decreases until 2017. In parallel, average term depth is largely maintained across subontologies, except again for GO:BP, where depth decreases until approximately 2011. Taken together, the increase in internal (non-leaf) nodes alongside a reduction in average depth indicates that growth in GO:BP has primarily occurred through lateral expansion. In other words, new internal terms tend to branch existing paths rather than extend them linearly away from the root, resulting in a “broader” rather than “deeper” ontology.

The most pronounced structural changes occur at the first layer of the ontology, which comprises the most general and least specific functional terms. This layer exhibits substantial temporal variability, with notable reorganizations in GO:MF and GO:CC around 2018. Such changes are striking, as this top-level partitioning of functional space might be expected to be the most consensual and therefore the most stable. Alterations at this level reflect major conceptual revisions in how core biological functions are organized within GO.

These structural changes observed between 2017 and 2019 are consistent with findings reported by (Valverde *et al*. 2025). Although their analysis focused primarily on the network properties of GO, they also identified a “tipping point” around 2017, which they interpreted as a major reorganization of the ontology aimed at improving conceptual clarity, in line with principles promoted by the Common Anatomy Reference Ontology (CARO). Our results provide complementary evidence for this restructuring across multiple structural and temporal dimension

GO annotations also show a general increase over time across all annotation resources analyzed. The volume, composition, and evidence profile of annotations contributed by different member databases reflect the depth and focus of experimental research conducted on their respective model organisms. Well-studied systems such as *Saccharomyces cerevisiae* and mouse (*Mus musculus*) exhibit dense and diverse annotation sets, driven by decades of targeted experimental work and sustained manual curation. Consequently, temporal trends in annotation content provide an indirect proxy for research intensity and knowledge accumulation, in addition to changes in annotation practices and pipelines.

The GOA dataset, maintained primarily by UniProt, occupies a distinct position within the GO annotation ecosystem due to its broad taxonomic coverage and its integration of extensive automated pipelines with selective manual curation. GOA provides annotations for a wide range of species, including many not covered by dedicated model organism databases, resulting in large annotation volumes dominated by electronically inferred annotations. In contrast, such automated annotations are less prevalent in organism-specific databases, where dedicated curators play a central role. At the same time, GOA incorporates high-quality manually curated annotations from Swiss-Prot entries, which are regularly reviewed and updated within UniProt workflows. This combination leads to annotation dynamics that differ substantially from those of organism-specific databases, and temporal changes in GOA annotation counts often reflect updates to UniProt pipelines, mapping resources, or quality-control procedures rather than changes in organism-specific experimental knowledge alone.

Overall, our analysis portrays GO as a dynamic—yet increasingly mature—resource, evolving in terms of its terms, the biological concepts they capture, and the complex network of relationships connecting them. These changes mirror, in part, the shifting priorities and conceptual frameworks of the biological research community. Structural modifications are most pronounced at the top levels of the ontology, reflecting deep conceptual revisions. The convergence of multiple indicators around a tipping point in approximately 2017, followed by increased stability thereafter, suggests that GO has reached a level of maturity in which future changes are likely to be more incremental.

The temporal evolution of GO, both in its structure and its associated annotations, has important implications for the development and maintenance of FAIR-compliant bioinformatics tools (Wilkinson *et al*. 2016). Changes in term definitions, relationships, and annotation coverage directly affect reproducibility, interoperability, and reusability of computational analyses. FAIR-aligned systems must therefore support explicit tracking of ontology and annotation versions, provide transparent provenance of data processing, and be updated regularly. By accounting for the temporal dynamics of GO, bioinformatics tools can better integrate historical and current datasets, ensure consistent interpretation of results across releases, and maintain long-term reproducibility.

Finally, for end users—particularly those performing enrichment or functional interpretation analyses—it is essential to be aware of the temporal dependence of GO-based results. Past analyses should be interpreted in the context of the GO and annotation versions used, and best practice dictates that tools, ontology versions, and annotation releases be explicitly reported to support transparency and reproducibility.

## Materials and methods

### GO ontology datasets

We retrieved GO terms and relationships from OBO files (go.obo), which provide a complete description of each GO term, including its identifier, name, definition, synonyms, and relationships to other terms within the ontology. The file also explicitly labels deprecated (“obsolete”) terms. We obtained go.obo files from the GO Archive (https://release.geneontology.org/) for each year from 2004—the earliest year available in the GO archive—through 2024.

### GO annotation datasets

Gene annotations were downloaded from the GO Archive (https://release.geneontology.org/). We downloaded annotation data from three resources representing different curation strategies and organismal scopes: Mouse Genome Informatics (MGI), providing annotations for *Mus musculu*s; Saccharomyces Genome Database (SGD), for *Saccharomyces cerevisiae*; and annotations from GOA-UniProt, which supplies GO annotations for proteins in the UniProt Knowledgebase. All annotation files include evidence codes describing the type of support for each annotation (except for GOA that does not provide IEA annotations prior to 2018).

Analogous to the ontology data, we downloaded yearly annotation releases for each database. Historical data were available from 2004 onward for MGI, SGD, and WormBase, and from 2010 onward for GOA-UniProt.

### Temporal analysis of GO terms

For each year, we compared the list of GO terms to that of the preceding year in order to identify newly introduced terms and newly obsoleted terms. This procedure yielded, for each year, the total number of terms, the number of new terms, and the number of newly obsoleted terms. All counts were stratified by GO subontology—Biological Process (BP), Molecular Function (MF), and Cellular Component (CC)—and normalized using Z-scores to facilitate comparison across subontologies and time periods.

We defined the lifespan of a GO term as the number of years between its first appearance and the year it was marked obsolete, or until 2024 for terms that remain active.

### Word enrichment analysis of GO term additions

To characterize the biological themes emphasized in GO at different time points, we performed a word enrichment analysis on newly added GO terms for each year. From the yearly OBO files, we extracted individual words appearing in term names and exact synonyms. For example, for GO:0000024 (“maltose biosynthesis”), with the exact synonym “malt sugar biosynthesis”, the extracted words are maltose, biosynthesis, malt, and sugar. In addition to single words, we also considered pairs of consecutive words (bigrams).

To identify words enriched in a given year, we applied the hypergeometric survival function test. For a given word of a particular year, we calculated the p-value of the null hypothesis (i.e. the word is not enriched that year but it presents the expected frequency). The resulting p-values quantify the statistical significance of word enrichments.

For each year, the 25 words with smallest p-values (with p-value < 0.001) were visualized using a word cloud representation, in which word size is proportional to the negative base-10 logarithm of the corresponding p-value.

### Ontology structure metrics

The depth of a GO term was computed as the length of the shortest path separating the term from the root node of its corresponding subontology: “biological process” (GO:0008150), “cellular component” (GO:0005575), or “molecular function” (GO:0003674).

Leaf terms were defined as terms with no descendants in the directed acyclic graph (DAG), representing the most specific functional concepts. First-layer terms were defined as those having a direct relationship with the root term of their respective subontology.

GO relationships were grouped into three categories for analysis: *is_a* relationships, *part_of* relationships, and other relationships, the latter encompassing all regulatory relationships, *occurs_in*, and *capable_of* relations.

### Evidence code analysis

Finally, we analyzed GO protein annotations retrieved from MGI, SGD, and GOA-UniProt together with their associated evidence codes grouped in six broad classes as defined by GO (Table S3), enabling the stratification of annotation dynamics by evidence type across time and annotation resources.

## Funding

This work was supported by grant PID2022-140017OB-C22 (MC and FP), funded by MICIU/AEI/10.13039/501100011033 and by “ERDF/EU”, and by grant PRE2020-095361 (BP), funded by MICIU/AEI /10.13039/501100011033 and by FSE+.

## Acknowledge of AI use

The authors used ChatGPT to improve spelling, grammar, clarity, and readability during manuscript preparation. After using this tool, the authors carefully reviewed and edited the text and took full responsibility for the final content.

## Supplementary figures

**Figure S1.**
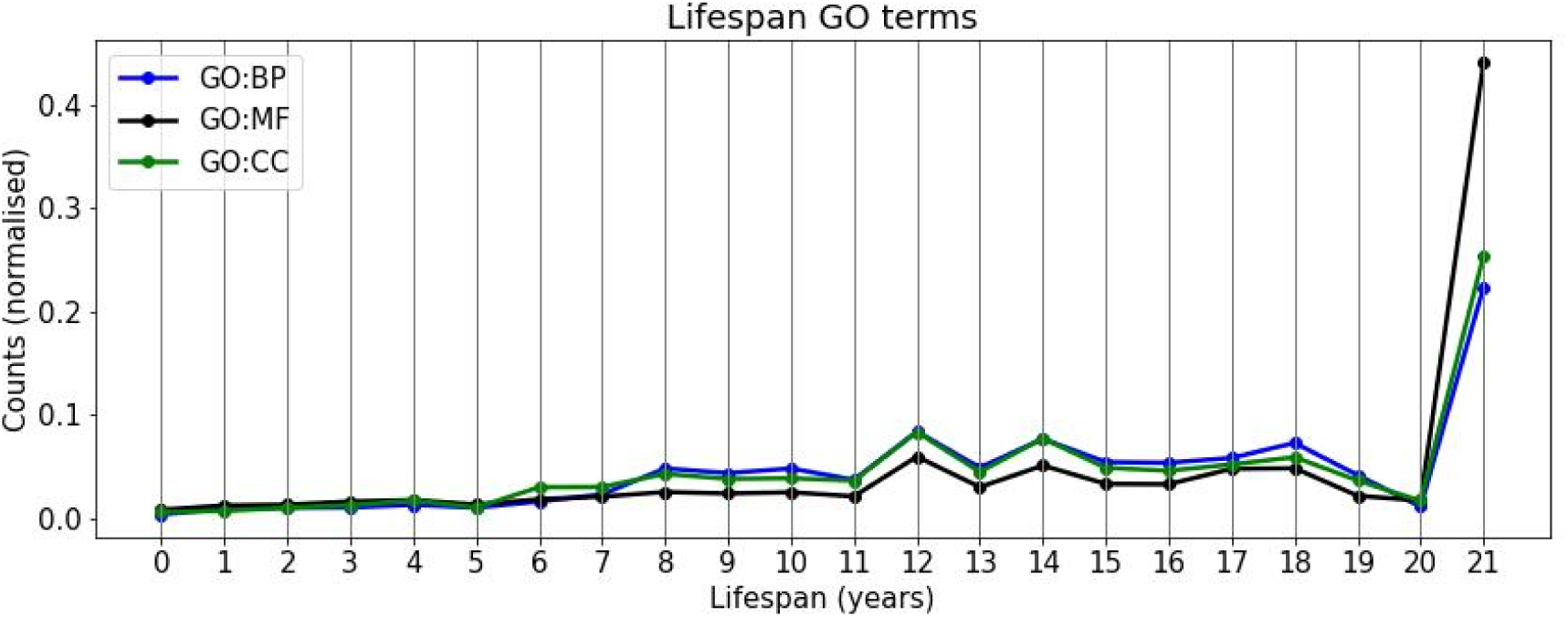
Lifespan of GO terms in the three subontologies.

**Figure S2:**
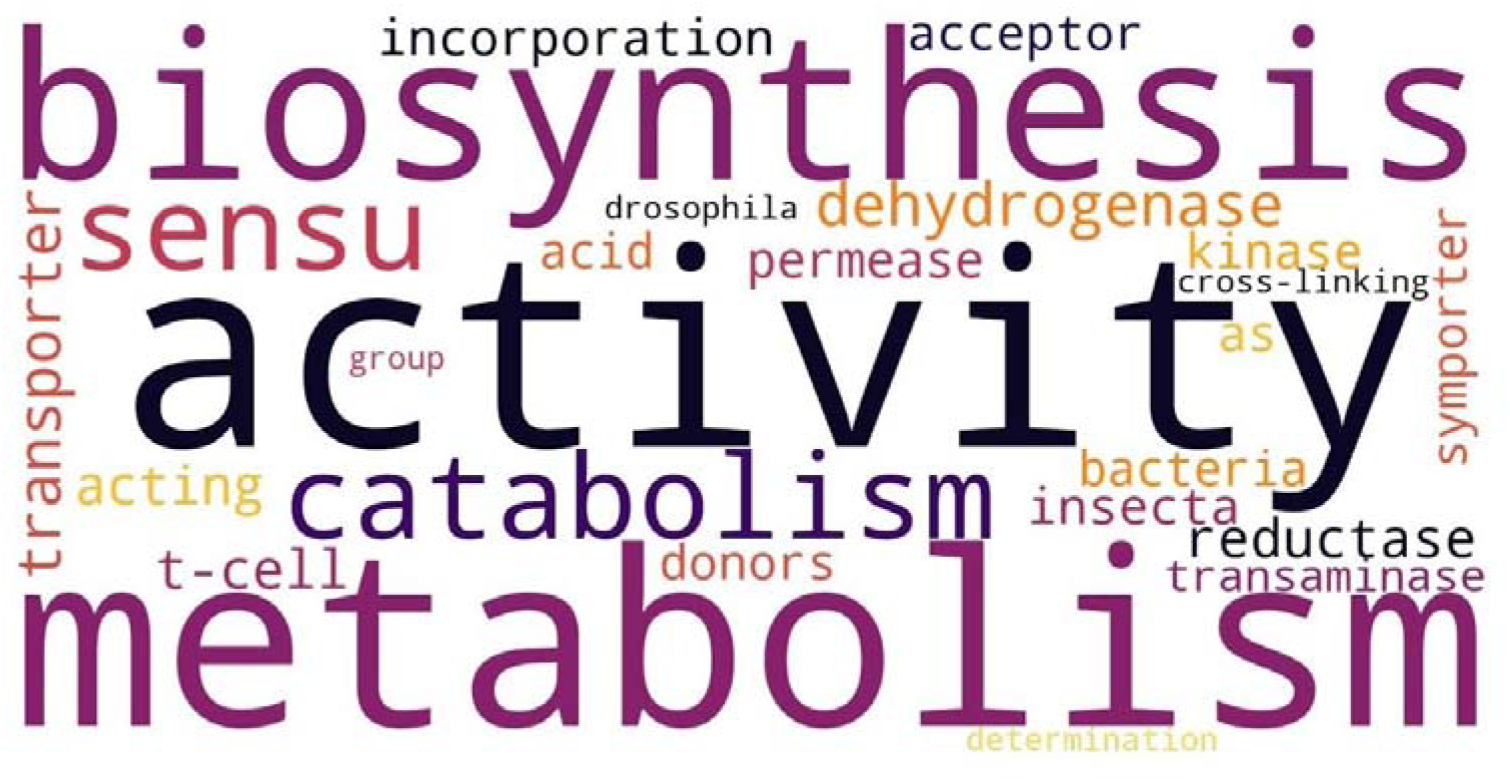
main words enriched in the names/synonyms of terms added to GO in 2004.

**Figure S3:**
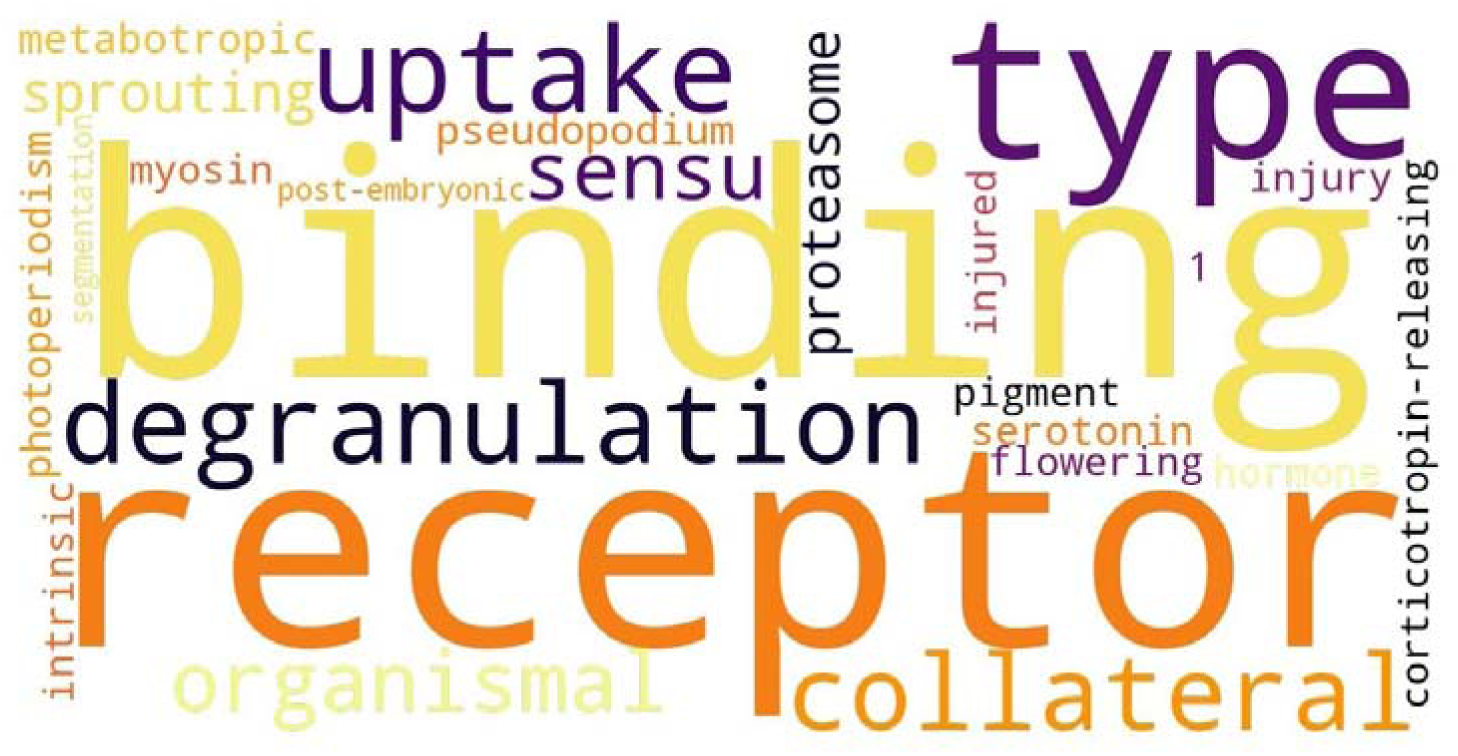
main words enriched in the names/synonyms of terms added to GO in 2005.

**Figure S4:**
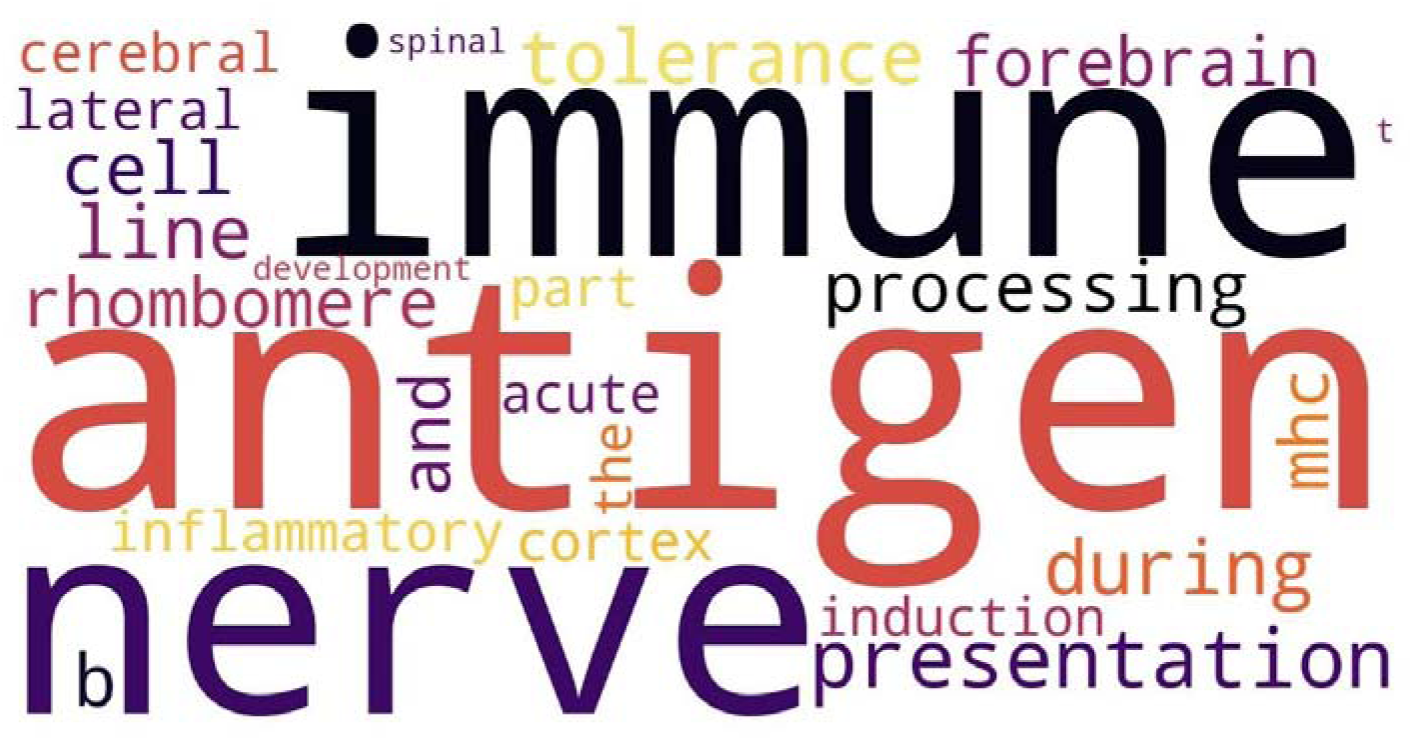
main words enriched in the names/synonyms of terms added to GO in 2006.

**Figure S5:**
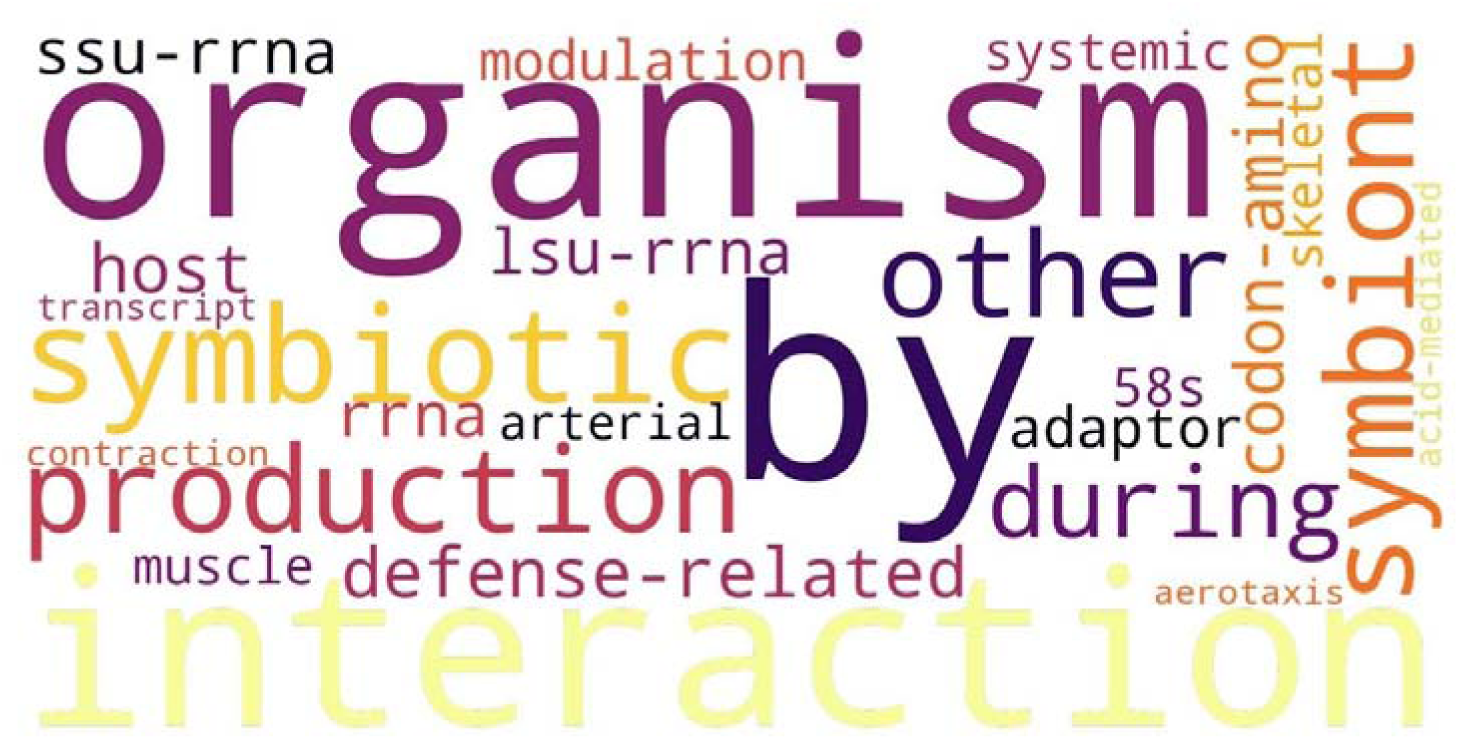
main words enriched in the names/synonyms of terms added to GO in 2007.

**Figure S6:**
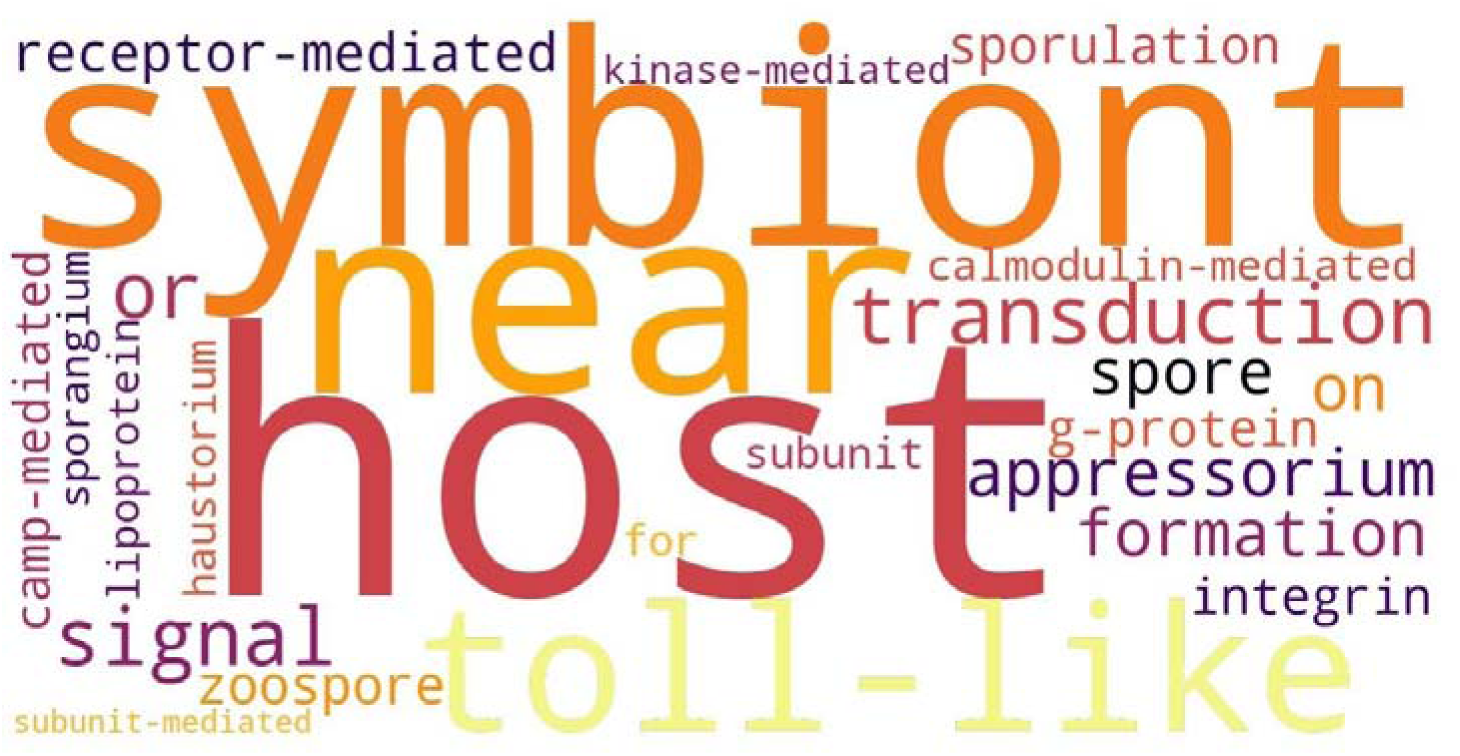
main words enriched in the names/synonyms of terms added to GO in 2008.

**Figure S7:**
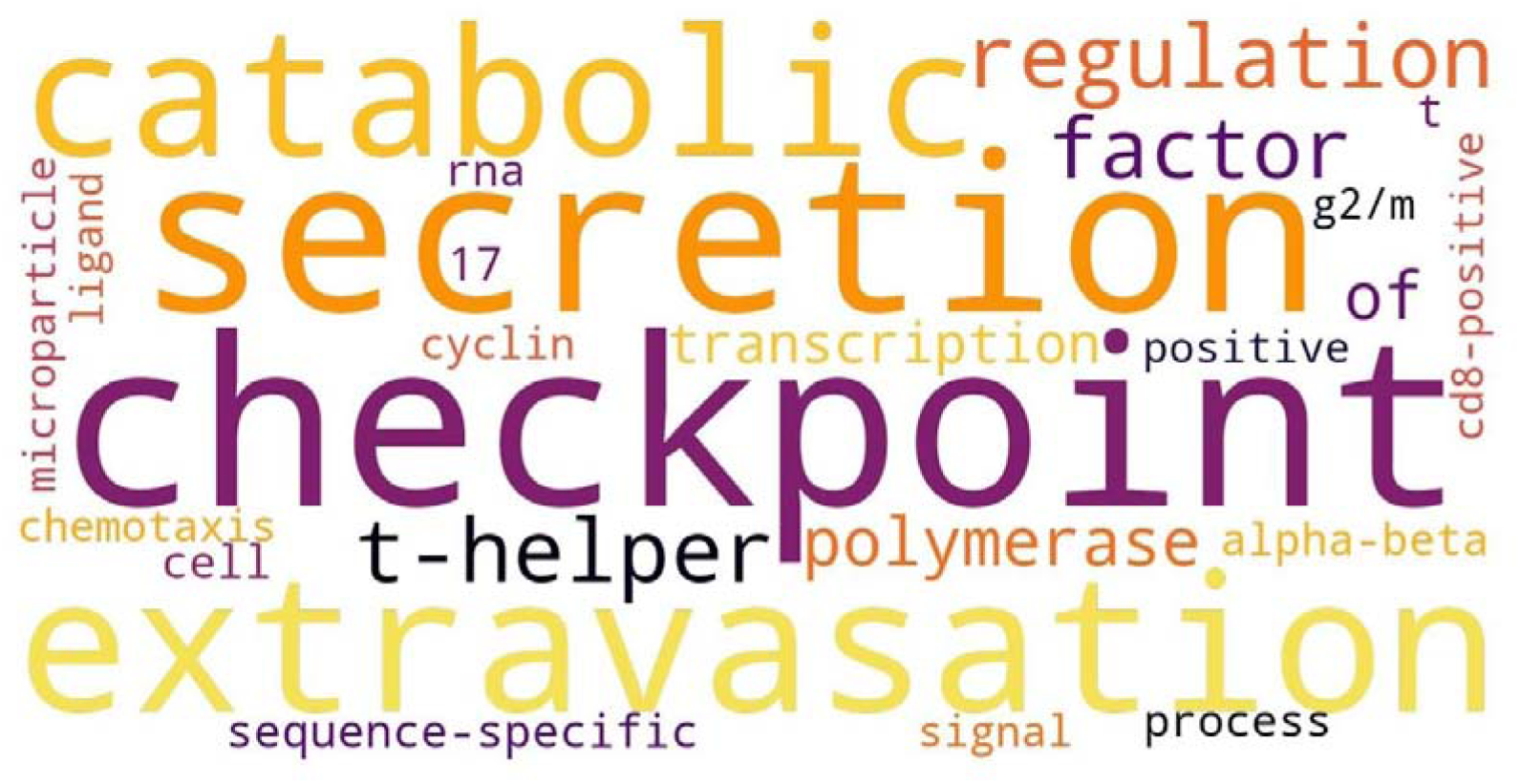
main words enriched in the names/synonyms of terms added to GO in 2011.

**Figure S8:**
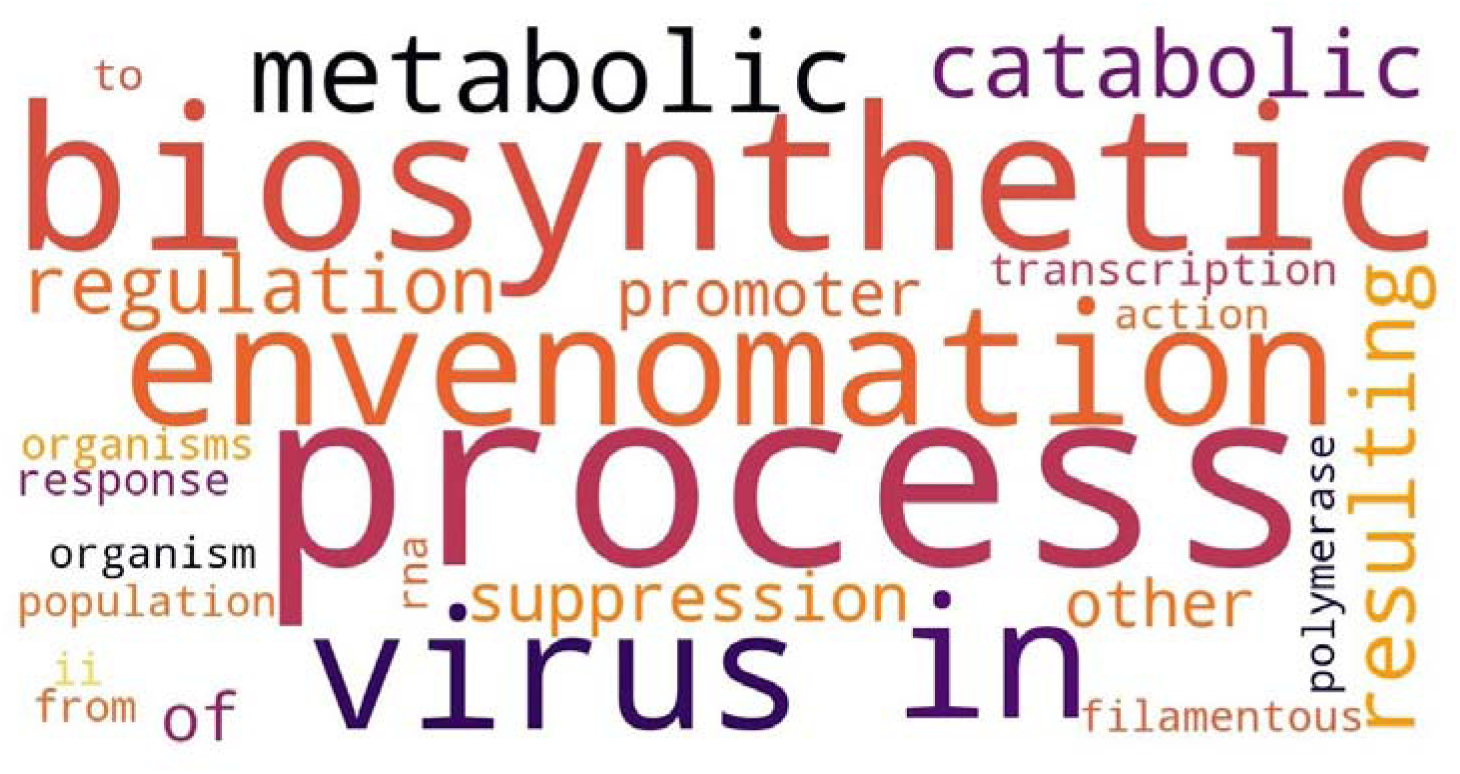
main words enriched in the names/synonyms of terms added to GO in 2012.

**Figure S9:**
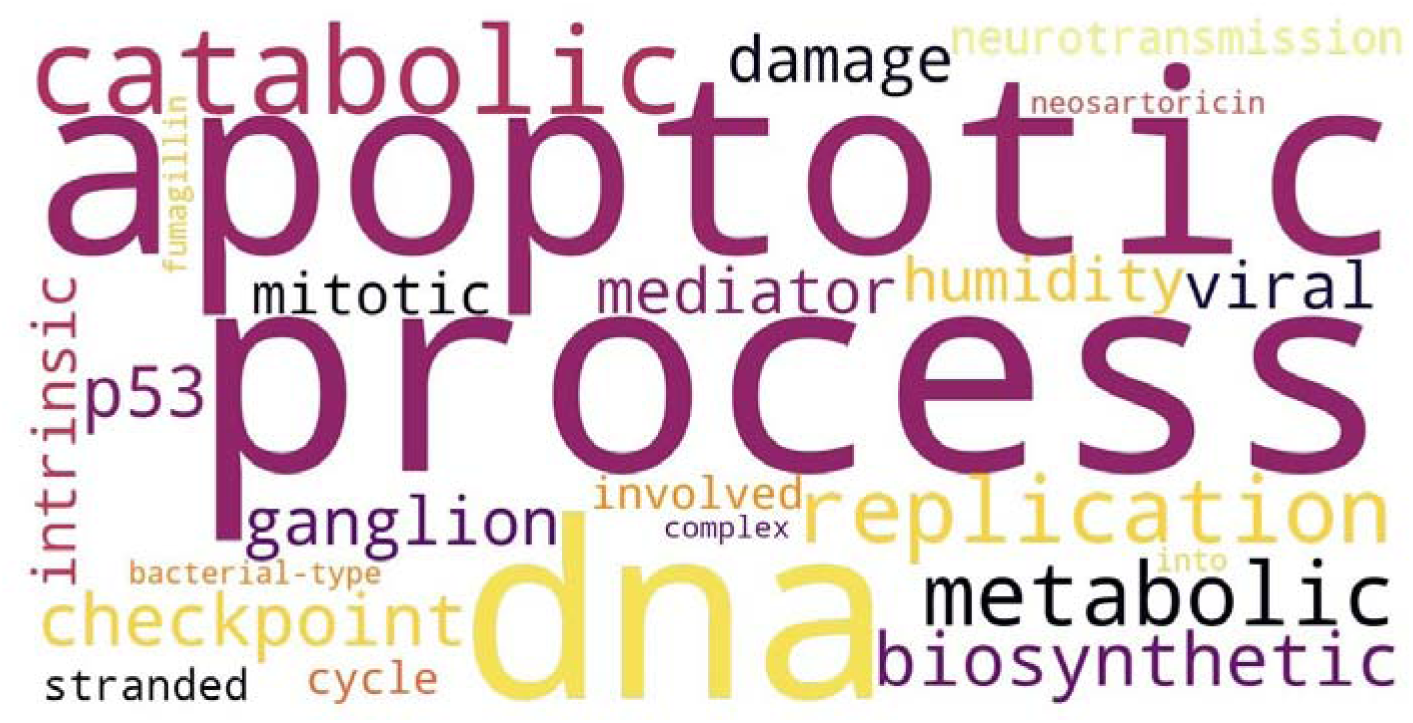
main words enriched in the names/synonyms of terms added to GO in 2013.

**Figure S10:**
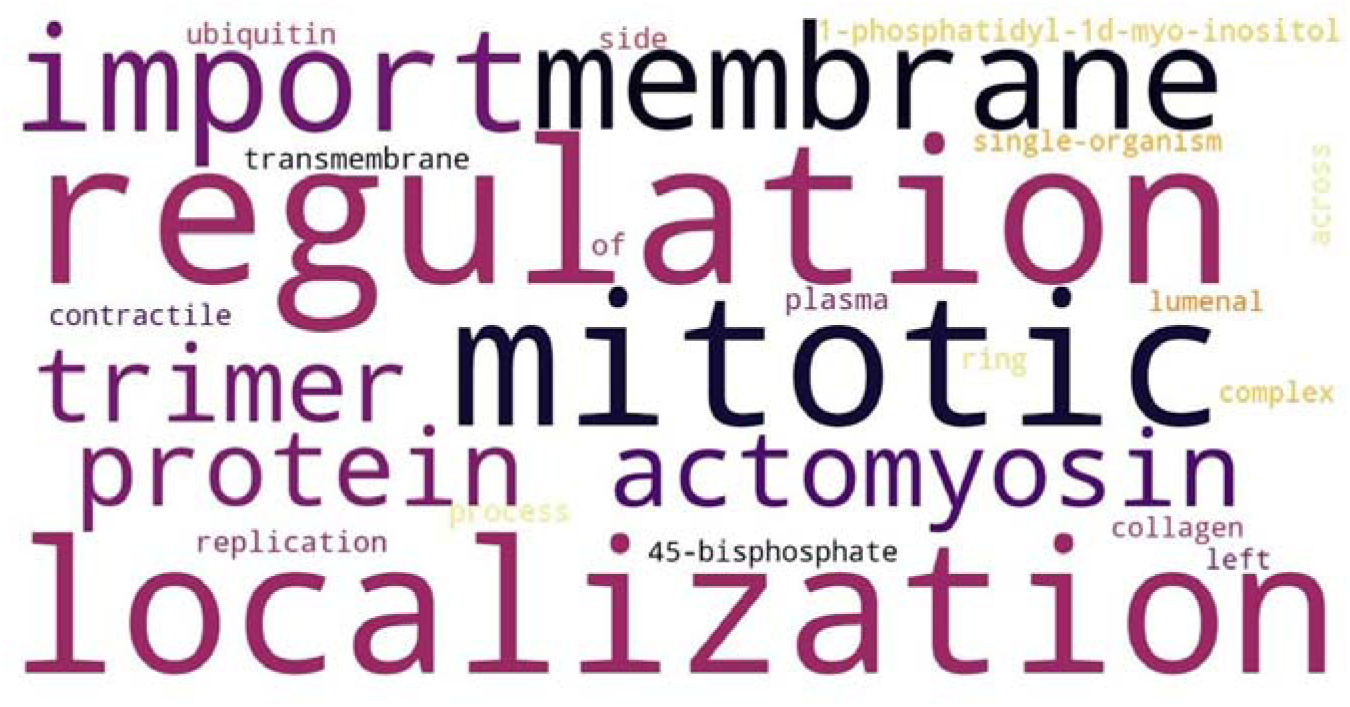
main words enriched in the names/synonyms of terms added to GO in 2014.

**Figure S11:**
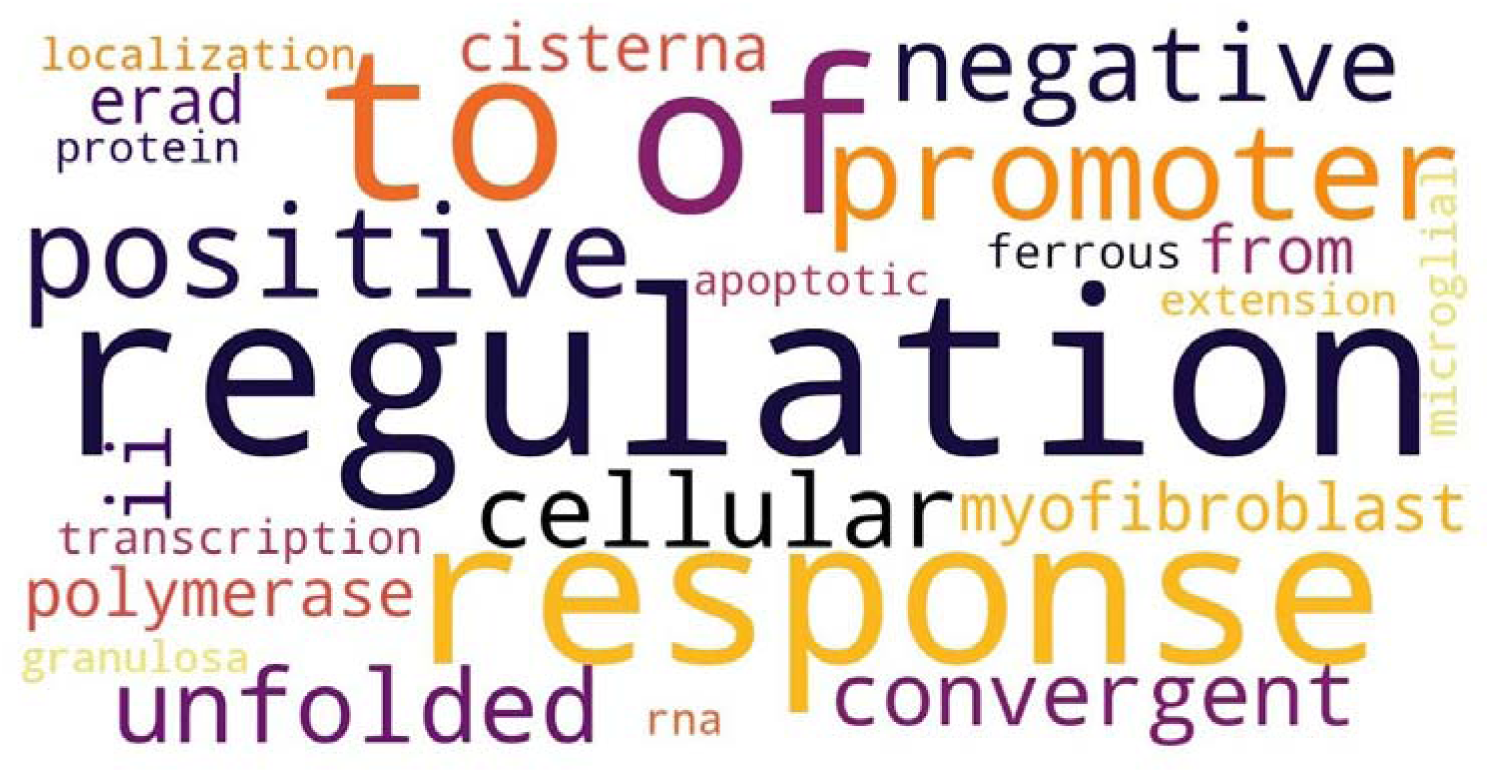
main words enriched in the names/synonyms of terms added to GO in 2015.

**Figure S12:**
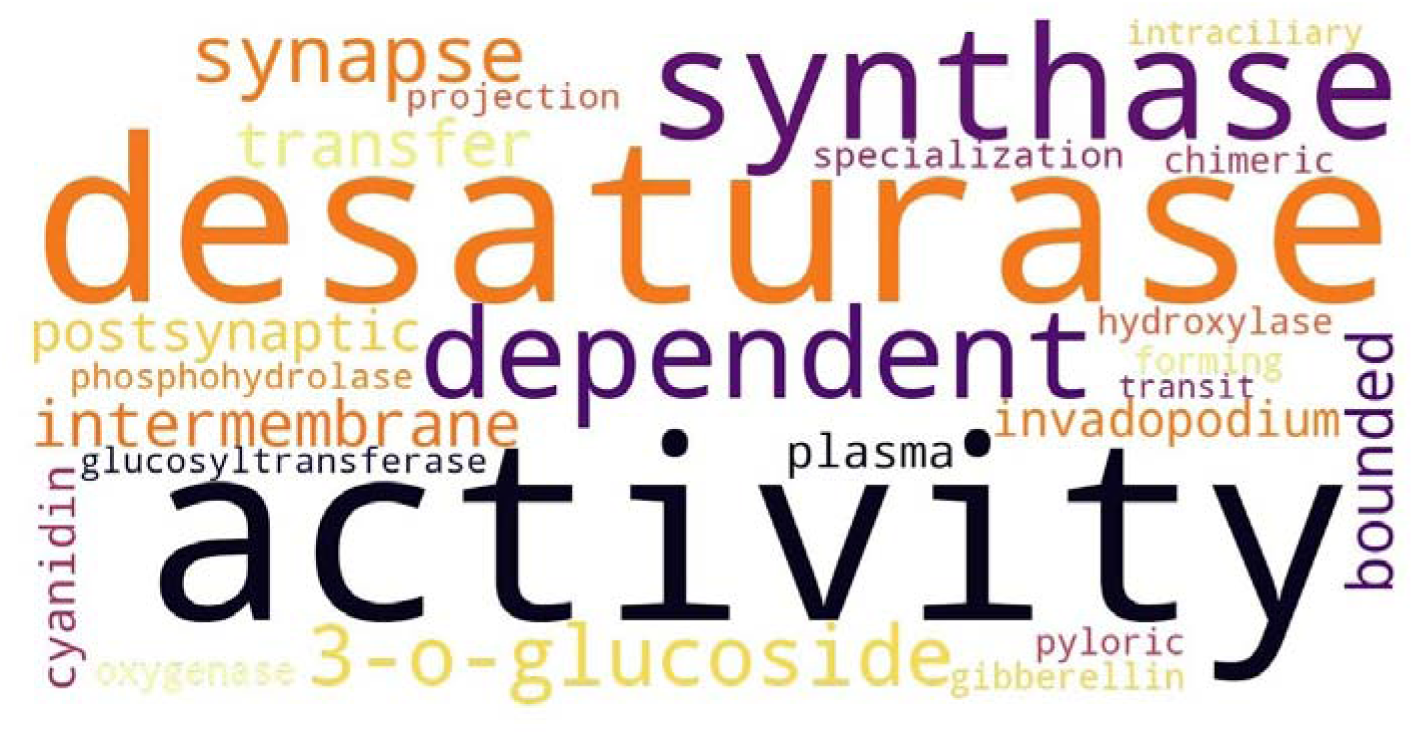
main words enriched in the names/synonyms of terms added to GO in 2017.

**Figure S13:**
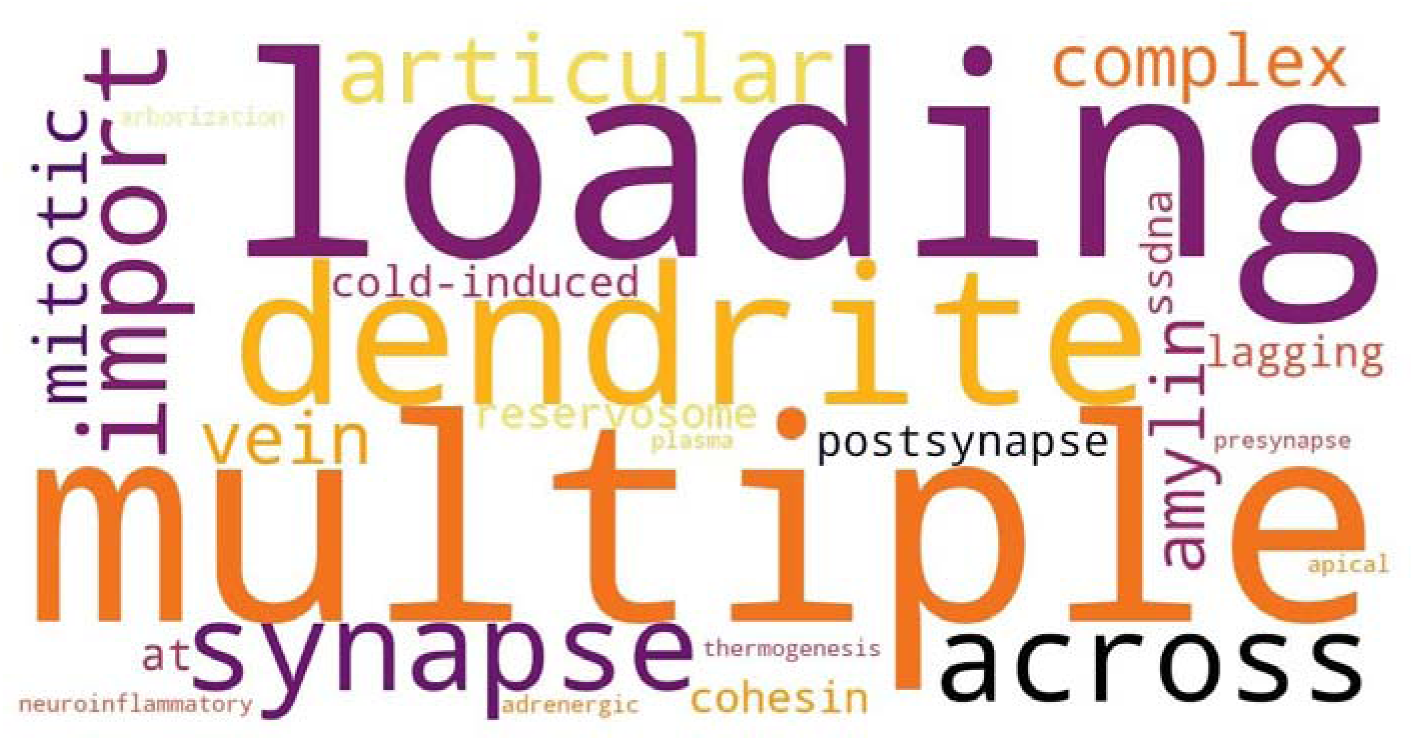
main words enriched in the names/synonyms of terms added to GO in 2018.

**Figure S14:**
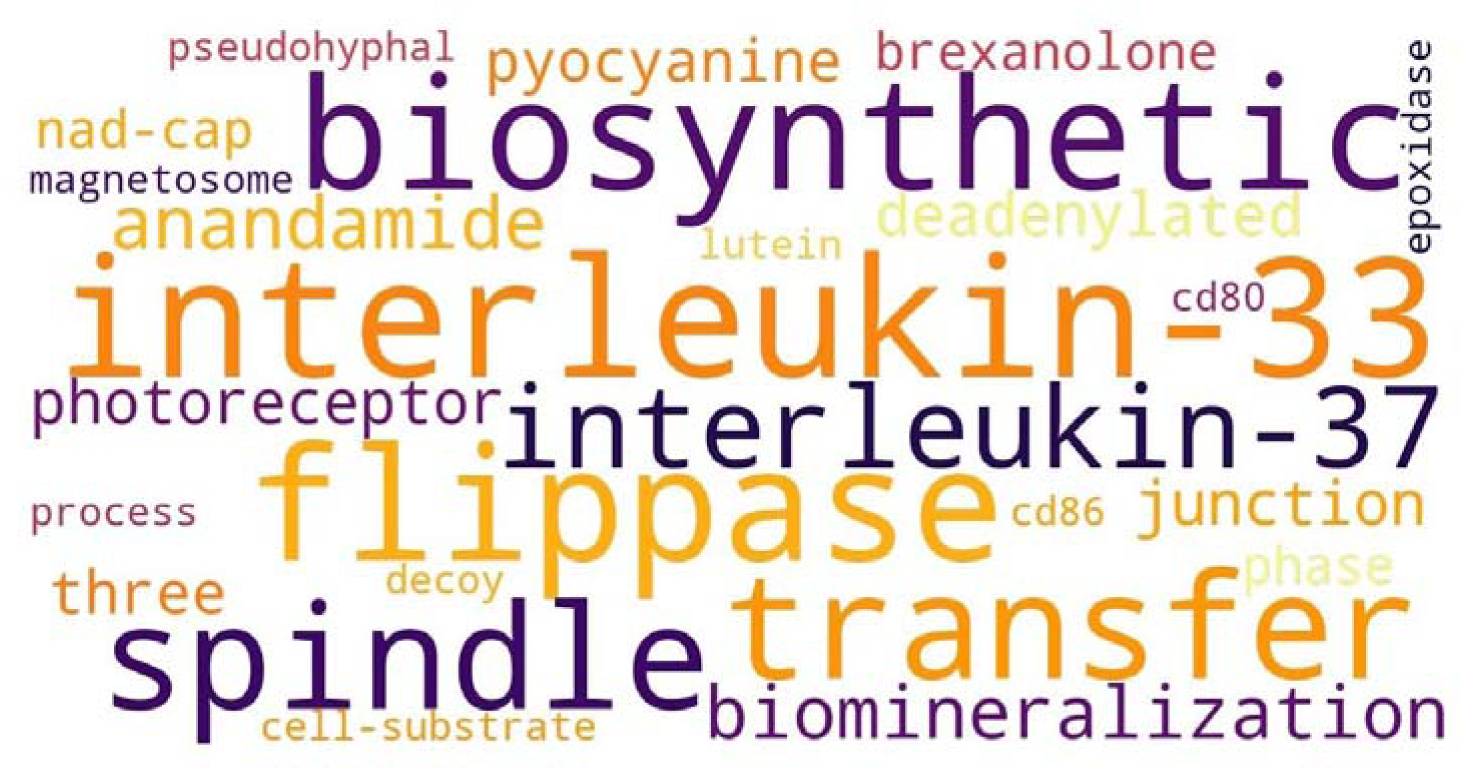
main words enriched in the names/synonyms of terms added to GO in 2019.

**Figure S15:**
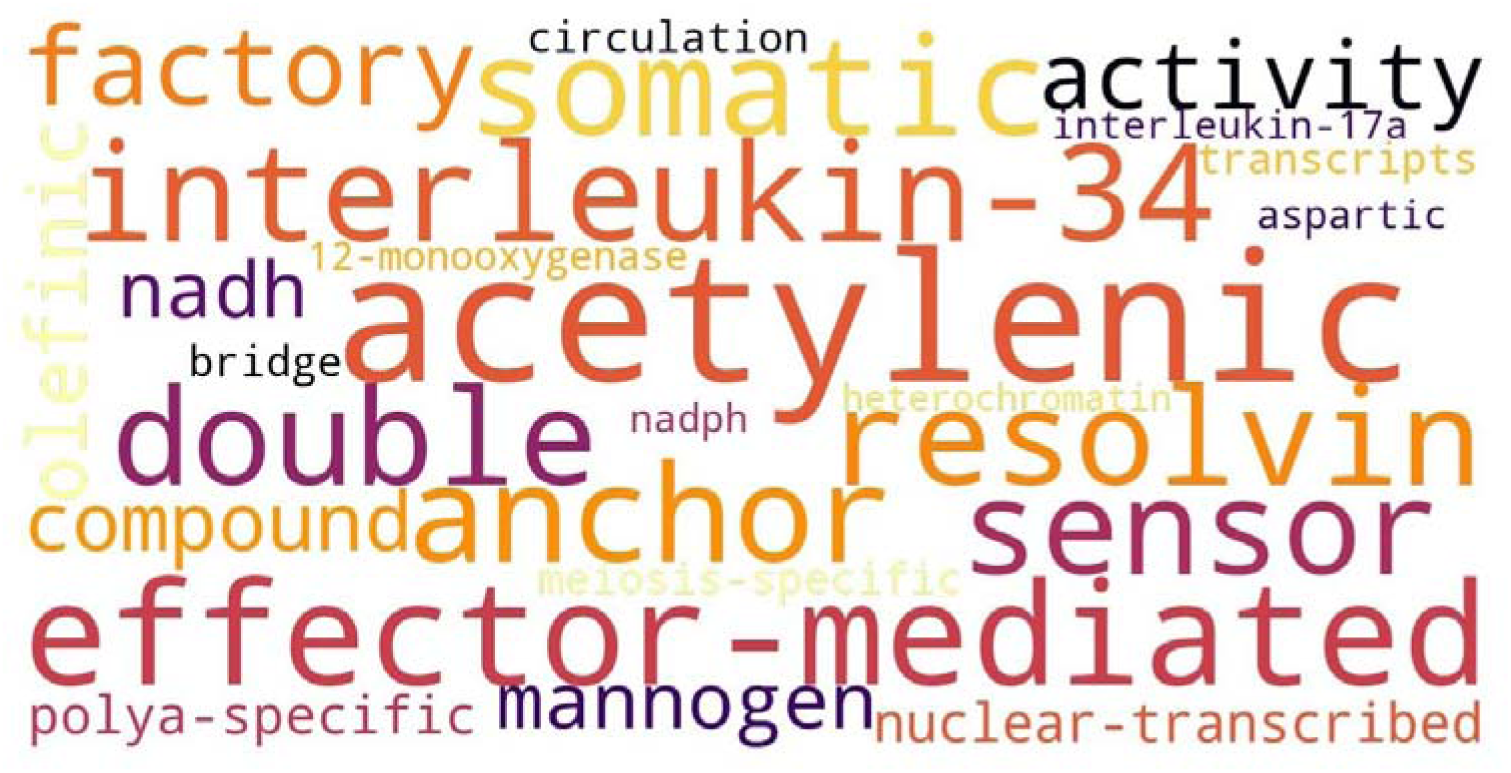
main words enriched in the names/synonyms of terms added to GO in 2020.

**Figure S16:**
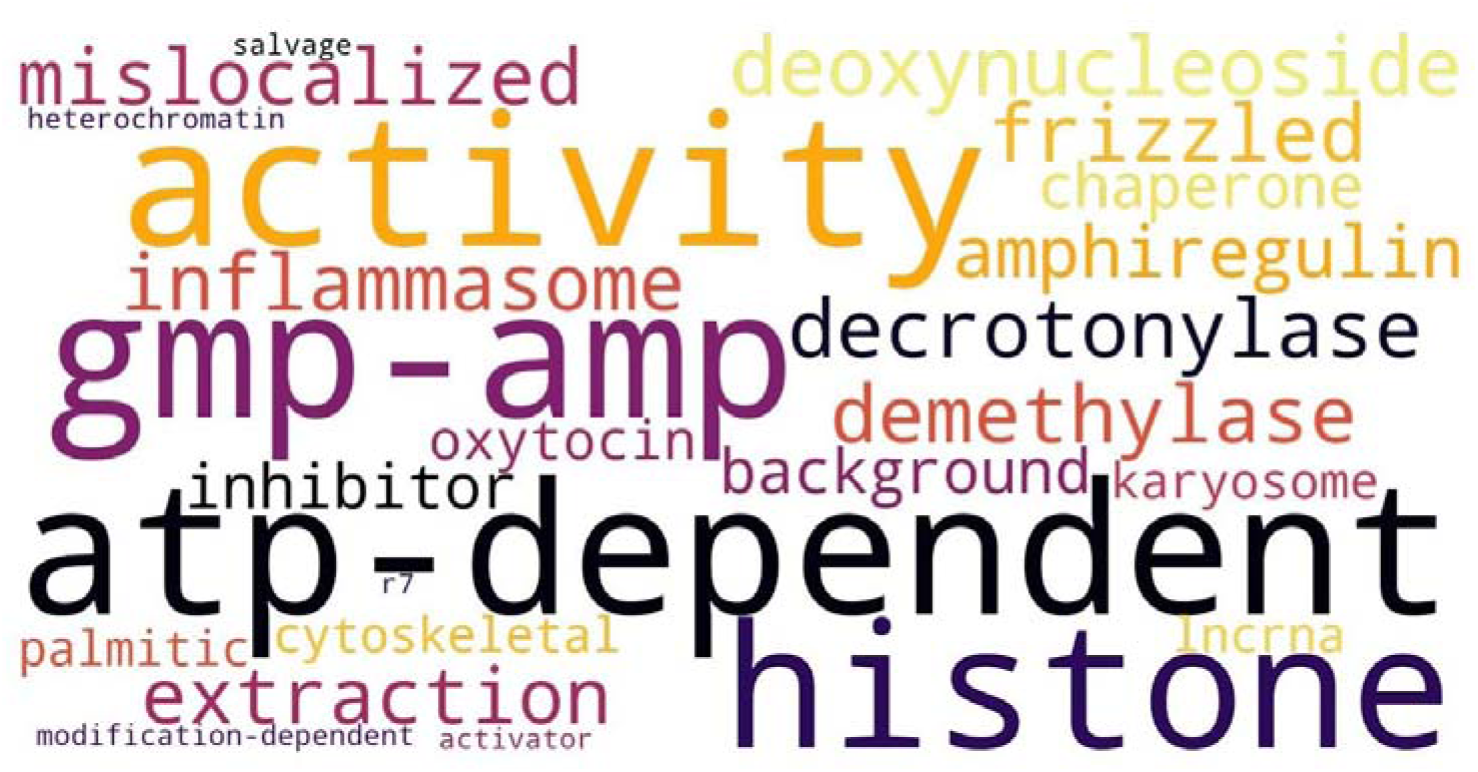
main words enriched in the names/synonyms of terms added to GO in 2021.

**Figure S17:**
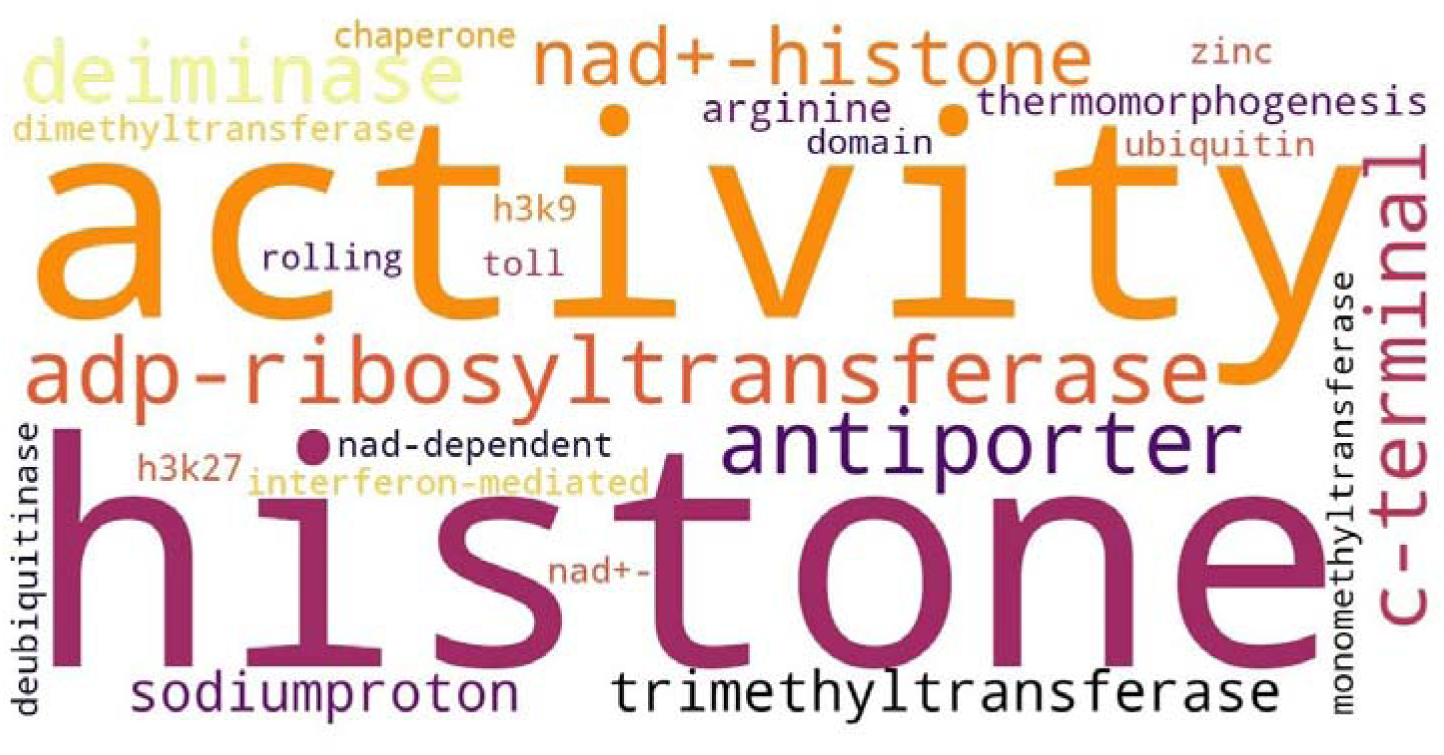
main words enriched in the names/synonyms of terms added to GO in 2022.

**Figure S18:**
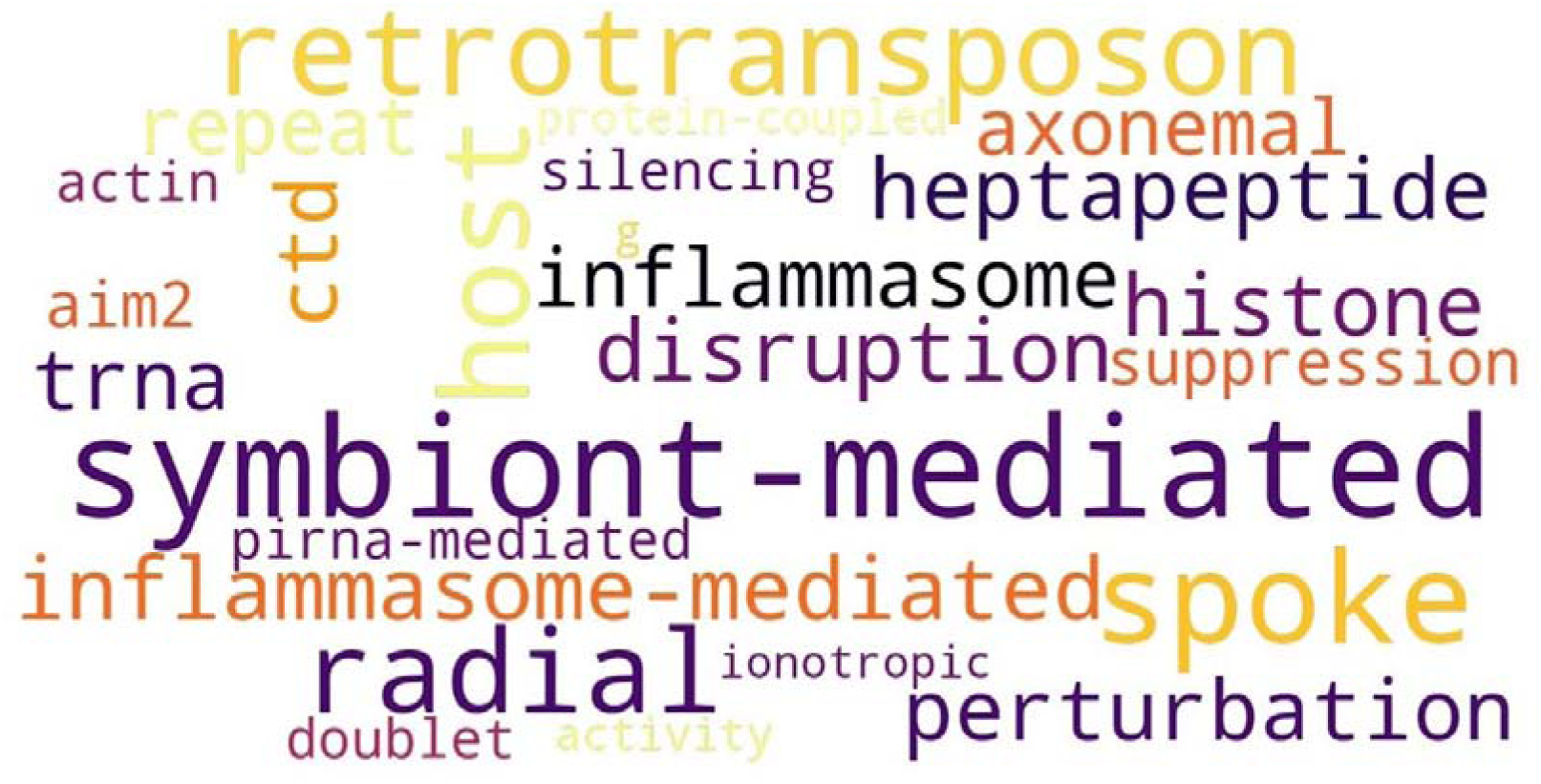
main words enriched in the names/synonyms of terms added to GO in 2023.

**Figure S19:**
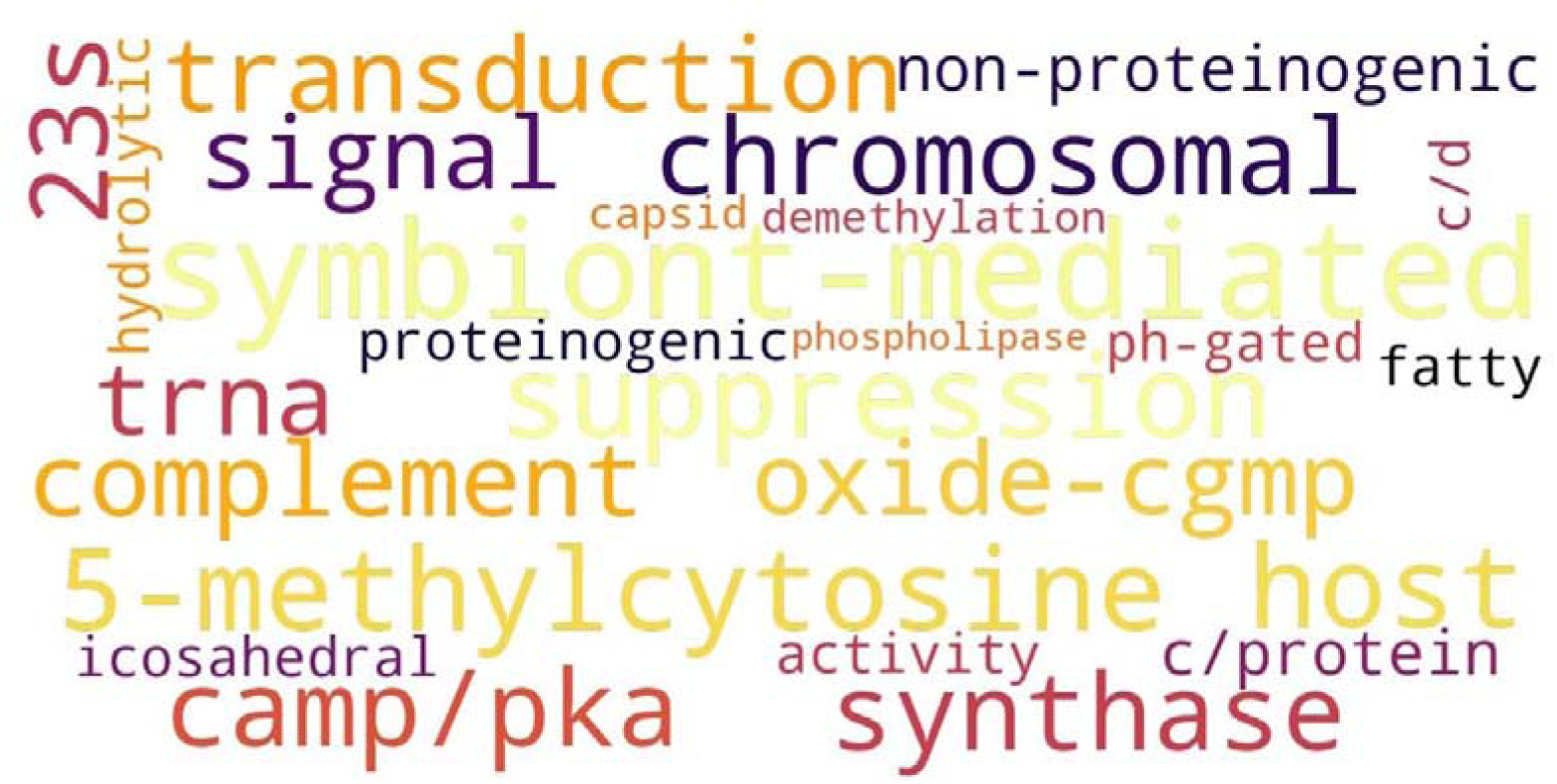
main words enriched in the names/synonyms of terms added to GO in 2024.

**Figure S20.**
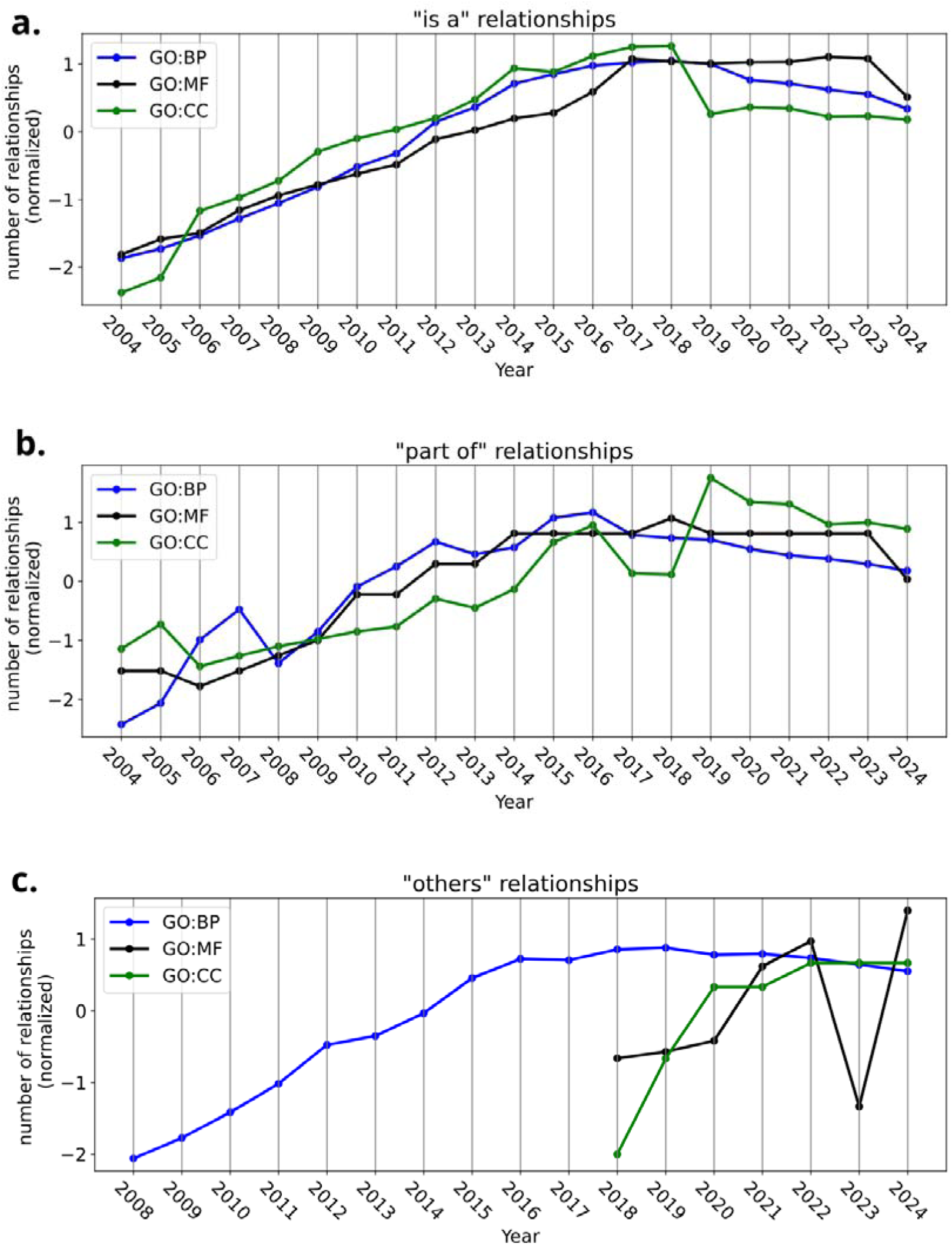
Evolution of number of relationships normalised. a) *is_a*, b) par_of, and c) others

**Figure S21.**
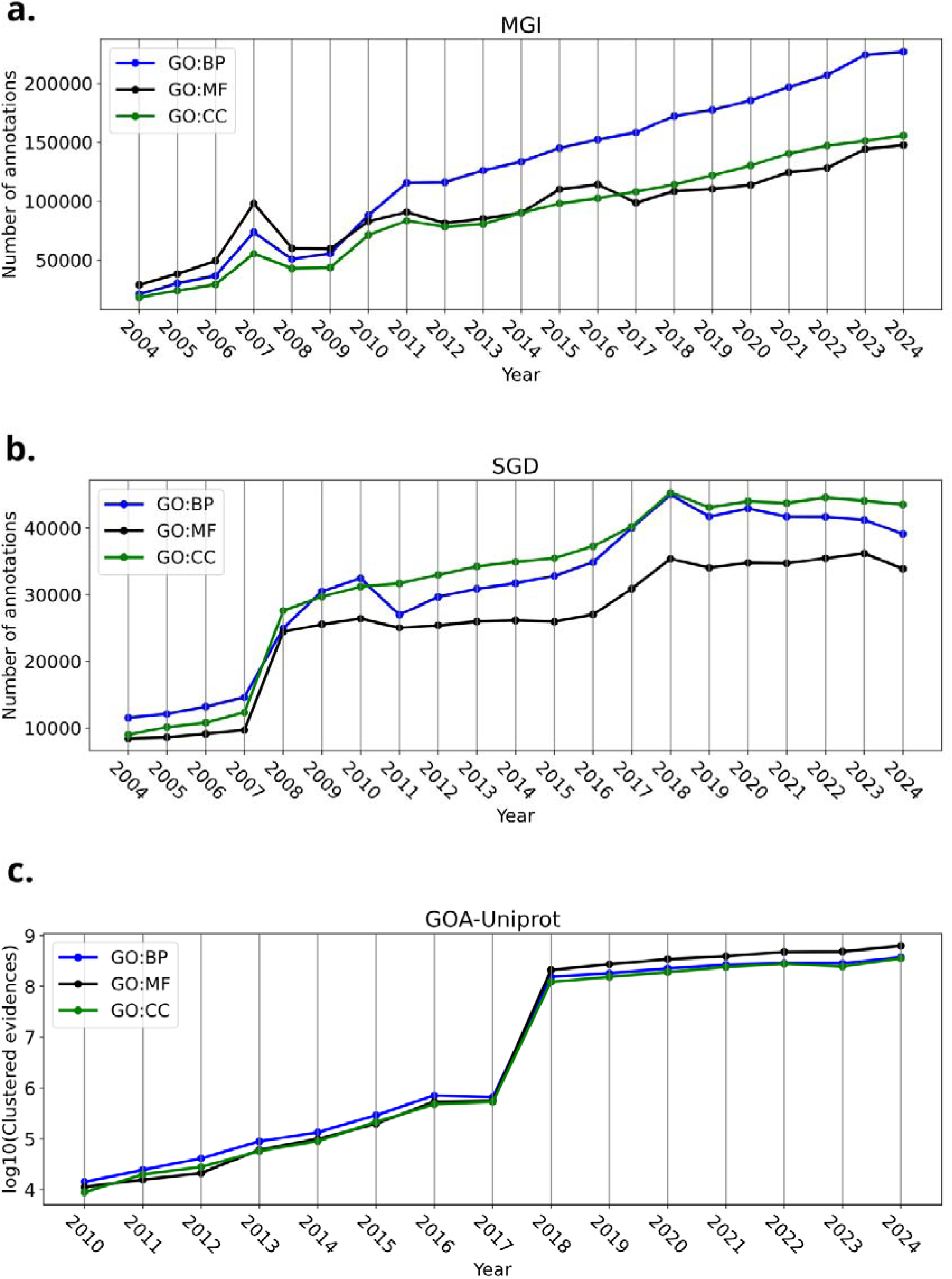
Number of GO annotations per subontology type in a) MGI (*M. musculus*), b) SGD (*S. cerevisiae*) and c) GOA-Uniprot (multi-species).

## Supplementary tables

**Table S1:** release dates for every go.obo dataset used.

| Year | Release date go.obo (DD/MM/YYYY) |
| --- | --- |
| 2004 | 01/12/2004 |
| 2005 | 01/12/2005 |
| 2006 | 01/12/2006 |
| 2007 | 01/12/2007 |
| 2008 | 01/12/2008 |
| 2009 | 01/12/2009 |
| 2010 | 01/12/2010 |
| 2011 | 01/12/2011 |
| 2012 | 01/12/2012 |
| 2013 | 01/12/2013 |
| 2014 | 01/12/2014 |
| 2015 | 01/12/2015 |
| 2016 | 01/12/2016 |
| 2017 | 01/12/2017 |
| 2018 | 01/12/2018 |
| 2019 | 09/12/2019 |
| 2020 | 08/12/2020 |
| 2021 | 15/12/2021 |
| 2022 | 04/12/2022 |
| 2023 | 15/11/2023 |
| 2024 | 03/11/2024 |

**Table S2:** release dates for the annotations datasets used.

| Year | Release date MGI, SGD and WormBase annotations datasets (DD/MM/YYYY) | Release date GOA-Uniprot annotations dataset (DD/MM/YYYY) |
| --- | --- | --- |
| 2004 | 01/04/2004 | None |
| 2005 | 01/04/2005 | None |
| 2006 | 01/04/2006 | None |
| 2007 | 01/04/2007 | None |
| 2008 | 01/04/2008 | None |
| 2009 | 01/04/2009 | None |
| 2010 | 01/04/2010 | 01/03/2010 |
| 2011 | 01/04/2011 | 01/03/2011 |
| 2012 | 01/04/2012 | 01/03/2012 |
| 2013 | 01/04/2013 | 01/03/2013 |
| 2014 | 01/04/2014 | 01/03/2014 |
| 2015 | 01/04/2015 | 01/03/2015 |
| 2016 | 01/04/2016 | 01/03/2016 |
| 2017 | 01/04/2017 | 01/01/2017 |
| 2018 | 04/02/2018 | 06/03/2018 |
| 2019 | 17/04/2019 | 10/03/2019 |
| 2020 | 23/04/2020 | 25/03/2020 |
| 2021 | 01/05/2021 | 02/01/2021 |
| 2022 | 22/03/2022 | 10/03/2022 |
| 2023 | 01/04/2023 | 06/03/2023 |
| 2024 | 28/03/2024 | 28/03/2024 |

**Table S3:** Evidence groups in GO and the evidence codes they contain. . This information comes from the GO webpage (Guide to GO Evidence Codes, 2025)

| Evidence group | Evidence codes |
| --- | --- |
| Experimental | Inferred from Experiment<br>Inferred from Direct Assay<br>Inferred from Physical Interaction<br>Inferred from Mutant Phenotype<br>Inferred from Genetic Interaction<br>Inferred from Expression Pattern<br>Inferred from High Throughput Experiment<br>Inferred from High Throughput Direct Assay<br>Inferred from High Throughput Mutant Phenotype<br>Inferred from High Throughput Genetic Interaction<br>Inferred from High Throughput Expression Pattern |
| Phylogeny | Inferred from Biological aspect of Ancestor<br>Inferred from Biological aspect of Descendant<br>Inferred from Key Residues<br>Inferred from Rapid Divergence |
| Computational | Inferred from Sequence or structural Similarity<br>Inferred from Sequence Orthology<br>Inferred from Sequence Alignment<br>Inferred from Sequence Model<br>Inferred from Genomic Context<br>Inferred from Reviewed Computational Analysis |
| Author | Traceable Author Statement<br>Non-traceable Author Statement |
| Curator | Inferred by Curator<br>No biological Data available |
| Inferred electronic annotation (IEA) | Inferred electronic annotation (IEA) |

## Notes

### Competing Interest Statement

The authors have declared no competing interest.

## Bibliography

Ashburner M, Ball CA, Blake JA et al. Gene ontology: tool for the unification of biology. The Gene Ontology Consortium. Nat Genet 2000;25(1):25–9.

Baldarelli RM, Smith CL, Ringwald M et al. Mouse Genome Informatics: an integrated knowledgebase system for the laboratory mouse. Genetics 2024;227(1):iyae031. 10.1093/genetics/iyae031.

Chen J, Goudey B, Geard N et al. Integration of background knowledge for automatic detection of inconsistencies in gene ontology annotation. Bioinformatics 2024;40(Supplement_1):i390–400. 10.1093/bioinformatics/btae246.

Clarke EL, Loguercio S, Good BM et al. A task-based approach for Gene Ontology evaluation. Journal of Biomedical Semantics 2013;4(1):S4. 10.1186/2041-1480-4-S1-S4.

Dameron O, Bettembourg C, Le Meur N. Measuring the evolution of ontology complexity: the gene ontology case study. PLoS One 2013;8(10):e75993. 10.1371/journal.pone.0075993.

Engel SR, Aleksander S, Nash RS et al. Saccharomyces Genome Database: advances in genome annotation, expanded biochemical pathways, and other key enhancements. Genetics 2025;229(3):iyae185. 10.1093/genetics/iyae185.

Foulger RE, Denny P, Hardy J et al. Using the Gene Ontology to Annotate Key Players in Parkinson’s Disease. Neuroinform 2016;14(3):297–304. 10.1007/s12021-015-9293-2.

Gene Ontology Consortium. The Gene Ontology knowledgebase in 2026. Nucleic Acids Res 2026;54(D1):D1779–92. 10.1093/nar/gkaf1292.

Groß A, Hartung M, Prüfer K et al. Impact of ontology evolution on functional analyses. Bioinformatics 2012;28(20):2671–7. 10.1093/bioinformatics/bts498.

Huntley RP, Sawford T, Mutowo-Meullenet P et al. The GOA database: gene Ontology annotation updates for 2015. Nucleic Acids Res 2015;43(Database issue):D1057–1063. 10.1093/nar/gku1113.

Paul M, Anand A, Pyne S. Impact of the Continuous Evolution of Gene Ontology on Similarity Measures. In: Deka B, Maji P, Mitra S et al. (eds), Pattern Recognition and Machine Intelligence. Cham: Springer International Publishing, 2019, 122–9. 10.1007/978-3-030-34872-4_14.

Pitarch B, Chagoyen M, Ranea JAG et al. A review on Gene Ontology evaluations. Database (Oxford) 2025;2025:baaf058. 10.1093/database/baaf058.

Reimand J, Isserlin R, Voisin V et al. Pathway enrichment analysis and visualization of omics data using g:Profiler, GSEA, Cytoscape and EnrichmentMap. Nat Protoc 2019;14(2):482–517. 10.1038/s41596-018-0103-9.

Sternberg PW, Van Auken K, Wang Q et al. WormBase 2024: status and transitioning to Alliance infrastructure. Genetics 2024;227(1):iyae050. 10.1093/genetics/iyae050.

Tomczak A, Mortensen JM, Winnenburg R et al. Interpretation of biological experiments changes with evolution of the Gene Ontology and its annotations. Sci Rep 2018;8(1):5115. 10.1038/s41598-018-23395-2.

Valverde S, Vidiella B, Martínez-Redondo GI et al. Structural Changes in Gene Ontology Reveal Modular and Complex Representations of Biological Function. Mol Biol Evol 2025;42(6):msaf148. 10.1093/molbev/msaf148.

Wilkinson MD, Dumontier M, Aalbersberg IJJ et al. The FAIR Guiding Principles for scientific data management and stewardship. Sci Data 2016;3:160018. 10.1038/sdata.2016.18.

